# Renal tubule Cpt1a overexpression protects from kidney fibrosis by restoring mitochondrial homeostasis

**DOI:** 10.1101/2020.02.18.952440

**Authors:** Verónica Miguel, Jessica Tituaña, J.Ignacio Herrero, Laura Herrero, Dolors Serra, Paula Cuevas, Coral Barbas, Diego Rodríguez Puyol, Laura Márquez-Expósito, Marta Ruiz-Ortega, Carolina Castillo, Xin Sheng, Katalin Susztak, Miguel Ruiz-Canela, Jordi Salas-Salvadó, Miguel A. Martínez González, Sagrario Ortega, Ricardo Ramos, Santiago Lamas

## Abstract

Chronic kidney disease (CKD) remains a major epidemiological, clinical and biomedical challenge. During CKD, renal tubular epithelial cells (TECs) suffer a persistent inflammatory and profibrotic response. Fatty acid oxidation (FAO), the main source of energy for TECs, is reduced in kidney fibrosis and contributes to its pathogenesis. To determine if FAO gain-of-function (FAO-GOF) could protect from fibrosis, we generated a conditional transgenic mouse model with overexpression of the fatty acid shuttling enzyme carnitine palmitoyl-transferase 1 A (CPT1A) in TECs. *Cpt1a* knock-in (CPT1A KI) mice subjected to three different models of renal fibrosis (unilateral ureteral obstruction, folic acid nephropathy-FAN and adenine induced nephrotoxicity) exhibited decreased expression of fibrotic markers, a blunted pro-inflammatory response and reduced epithelial cell damage and macrophage influx. Protection from fibrosis was also observed when Cpt1a overexpression was induced after FAN. FAO-GOF restituted oxidative metabolism and mitochondrial number and enhanced bioenergetics increasing palmitate oxidation and ATP levels, changes also recapitulated in TECs exposed to profibrotic stimuli. Studies in patients evidenced decreased CPT1 levels and increased accumulation of short and middle chain acyl-carnitines, reflecting impaired FAO in human CKD. We propose that strategies based on FAO-GOF may constitute powerful alternatives to combat fibrosis inherent to CKD.

**Graphical abstract:** Overexpression of CPT1A impedes tubular epithelial renal cell dedifferentiation and a pro-fibrotic phenotype by restoring mitochondrial metabolic bioenergetics.

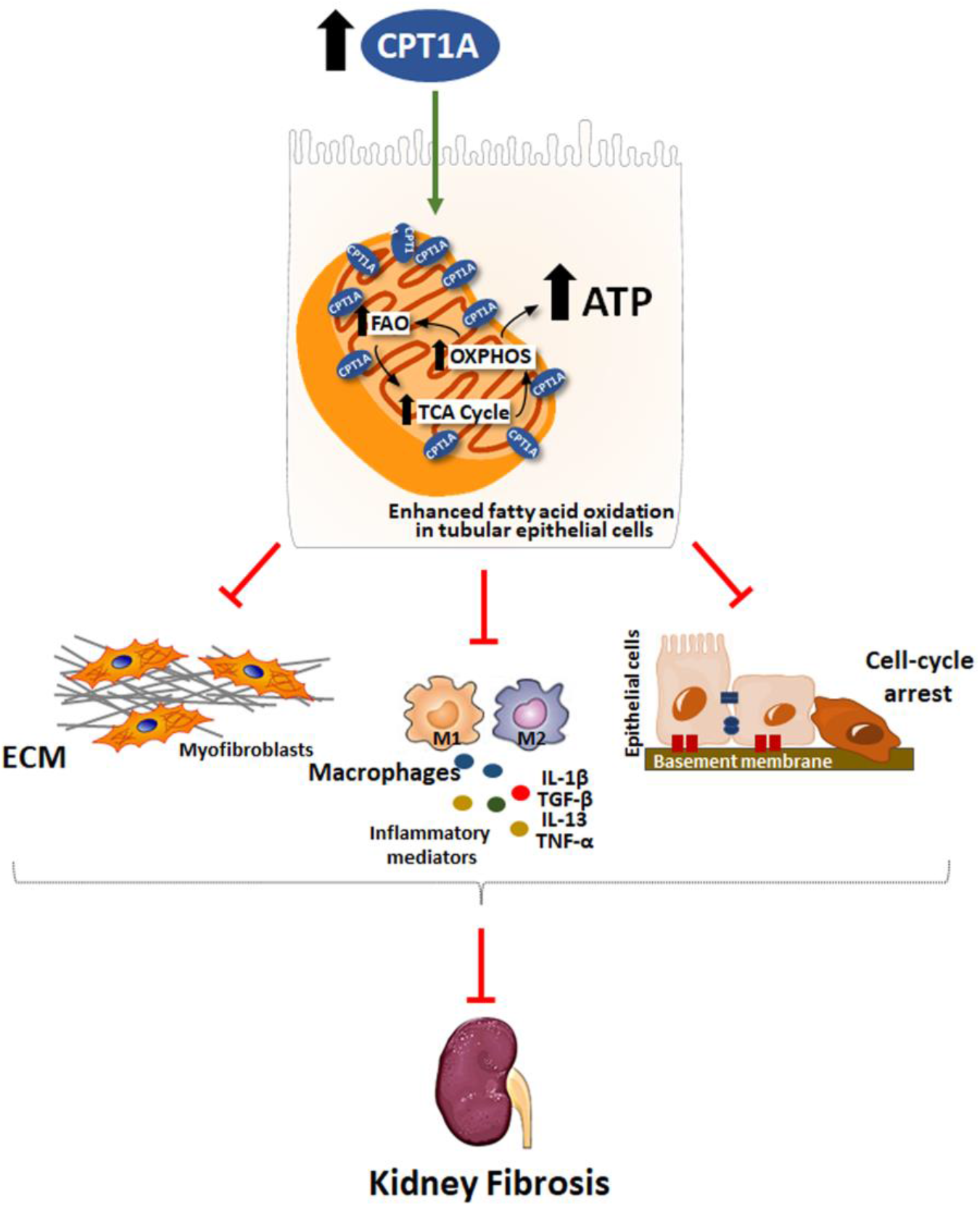

## Introduction

Organ fibrosis constitutes a significantly prevalent pathological entity associated to high morbidity and mortality and a major biomedical challenge. While it may affect any tissue of the human body, its presence in the kidney generally indicates unrelenting progression to chronic renal failure, a condition associated to reduced expectancy and quality of life. Hence, fibrosis is a convergent pathway for prevalent pathologies underlying chronic kidney disease (CKD) such as diabetes, hypertension or nephrosclerosis (1). Beyond proper blood pressure and glycemic controls, therapeutic options to revert or deter the progression of fibrosis are very limited. In the last few years, the understanding of major metabolic disturbances coexisting with kidney fibrosis have shed new light on the pathogenesis of fibrosis progression (2). Among these, a drastic reduction in fatty acid oxidation (FAO) appears to be critical for the global energy failure occurring in the tubulo-interstitial compartment, contributing to immune cell infiltration and kidney fibrosis (3, 4). Here we tested whether specific metabolic gain-of-function in FAO (FAO-GOF) in renal tubules was necessary and sufficient to counteract cellular and molecular changes associated to kidney fibrosis. Carnitine palmitoyl transferase 1A (CPT1A) is a rate-limiting and targetable enzyme in this pathway. We found that conditional overexpression of CPT1A resulted in significant mitigation of fibrosis and improvement of renal function in three different experimental models. This was related to enhanced mitochondrial mass, repaired architecture and bioenergetics recovery. The overexpression of CPT1A was also associated with a reduced inflammatory pattern and abrogation of TGF-beta-induced epithelial cell damage. In the folic acid nephropathy (FAN) model, pathological changes were markedly reverted when FAO-GOF was reinstated. Moreover, studies in patients with chronic renal failure showed that a reduction in the levels of CPT1A correlated with the degree of fibrosis. In addition, in a large cohort of diabetic patients with CKD, a specific profile of increased plasma acyl-carnitines was found, reinforcing the critical metabolic derangement of FAO associated to CKD.

## Results

### Overexpression of tubular CPT1A results in protection against kidney fibrosis

To test the *in vivo* relevance of CPT1A and FAO for renal fibrosis, a mouse model with conditional, inducible expression of *Cpt1a* was engineered as described in Methods. To ensure renal epithelial tissue specificity, these animals were crossed with mice expressing an optimized reverse tetracyclin-controlled transactivator (rtTA2s-M2), driven by the paired box 8 (Pax8) promoter, hereafter termed *Pax8-rtTA* mice. Extra-renal expression of Pax8 only occurs at the level of the thyroid without reported off-target effects (5). This results in tissue-specific expression of *Cpt1a* gene bearing TRE in proximal and distal tubules and collecting duct, which is evident two weeks after doxycycline administration (**Figure 1, A and B and Supplemental Figure 1, A and B**). We next characterized the renal specific overexpression of CPT1A in this newly generated genetic mouse model based on the doxycycline inducible transgenic system Tet-On. CPT1A KI mice presented a 10-fold increase in CPT1A mRNA level in the whole kidney tissue compared with WT mice (**Figure1C**). This was accompanied by a marked augment of CPT1A protein expression in the kidney. Of note, no differences were observed when the liver tissue was analyzed (**Figure 1, D and E**). Tubules were labeled by lotus tetragonolobus lectin (LTL), a marker for proximal tubules. Consistently, induced expression of CPT1A and GFP proteins co-localized in tubule segments as evaluated by immunohistochemistry (**Figure 1F**). Importantly, overexpressed CPT1A presented a mitochondrial localization pattern, which was observed by double immunostaining by using the ATP synthase beta-subunit (βF1) as a marker (**Figure 1G**). To assess the magnitude of CPT1A overexpression on FAO, we analyzed the capacity to oxidize radiolabeled palmitate by the renal tissue. Functionally, the increase in CPT1A protein in the renal epithelium increased the capacity of kidney tissue to oxidize ^14^C-palmitate as reflected in the levels of both ^14^C-palmitate-derived ^14^CO_2_ as well as in ^14^C-palmitate-derived acid-soluble products (**Figure 1H**).

**Figure 1.**
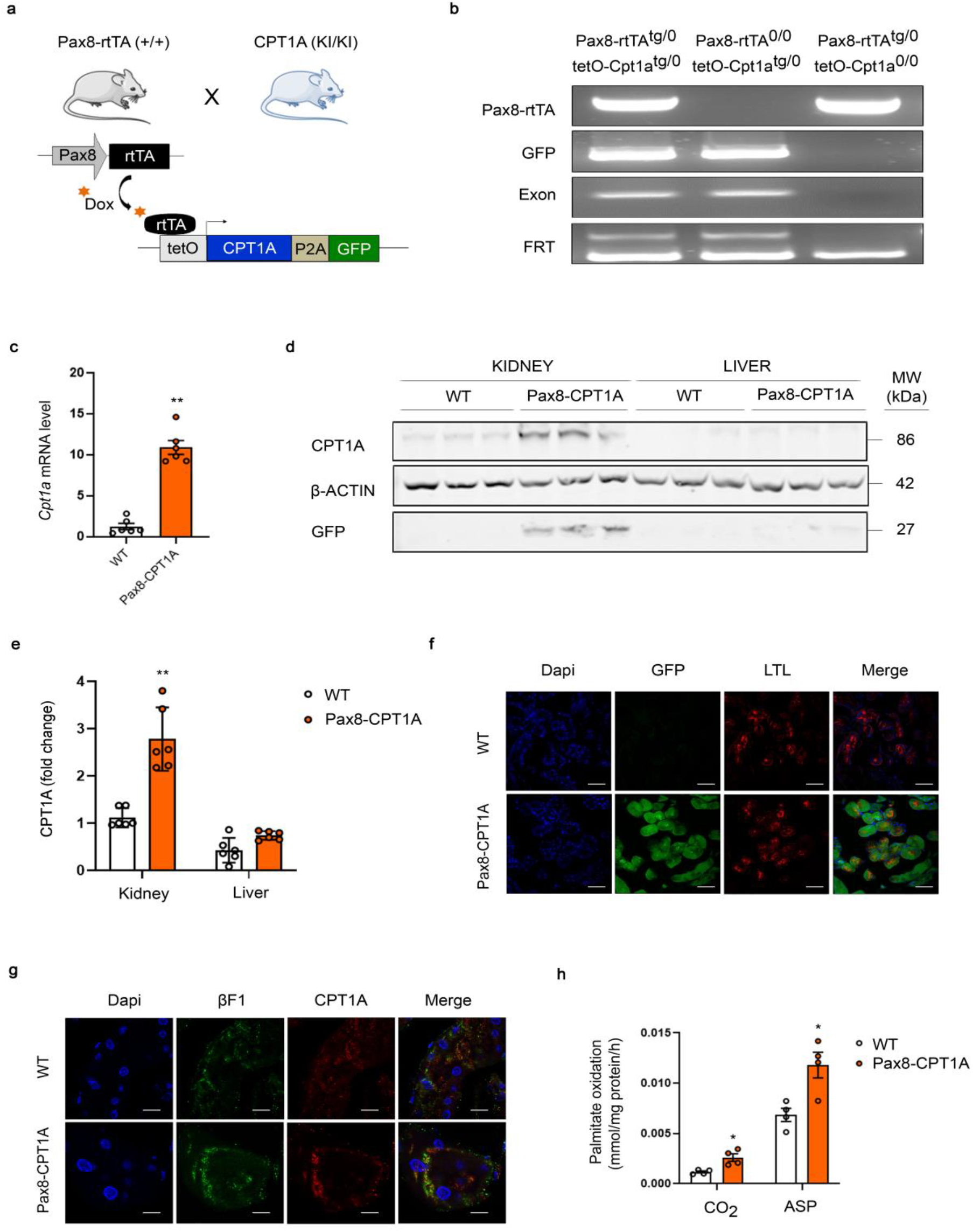
Characterization of Dox-inducible *Cpt1a* gene overexpression in the transgenic mouse model for inducible *Cpt1a* gene in renal epithelial cells. (**A**) Schematic depicting the strategy to generate mice with inducible renal tubular epithelial-specific overexpression of CPT1A. (see text for details). (**B**) PCR analysis of offspring genotypes from these crosses. Genomic DNA analysis by PCR for the Pax8-rtTA allele generates a 595-bp band (**Supplemental Table 1**). The GFP, Exon and FRT PCRs are described in **Supplemental Figure 1a**. (**C**) mRNA levels of *Cpt1a* gene was determined by qRT-PCR in total kidney tissue of mice treated with doxycycline for 3 weeks. Data represent the mean ± s.e.m (n = 6 mice). **P < 0.05 compared to kidneys from WT mice. (**D**) Immunoblots depicting protein levels of CPT1A and GFP in kidneys and livers of 3 individual mice per group. β-actin was used for normalization purposes. (**E**) Bar graphs represent the mean ± s.e.m of fold changes corresponding to densitometric analyses (n = 6 mice). **P < 0.01 compared to kidneys from WT mice. (**F**) Representative images of double immunofluorescence staining with the proximal tubular marker lotus tetragonolobus lectin (LTL), GFP, DAPI (nuclei) and merge of all three. (**G**) Staining with DAPI (nuclei), CPT1A and the mitochondrial marker ATP synthase beta-subunit (βF1). All panels show immunofluorescence images of kidneys from WT and Pax8-CPT1A mice after doxycycline administration. Scale bar: 100 μm (F and G). (**H**) Radiolabeled palmitate-derived CO_2_ and acid-soluble products (ASP) were determined after incubation of ^14^C-palmitate with kidney tissue from WT or Pax8-CPT1A mice after doxycycline treatment. Bar graphs represent the mean ± s.e.m (n = 4 mice). *P < 0.05 compared to kidneys from WT mice. Statistical significance between two independent groups was determined using non-parametric two-tailed Mann-Whitney test, while more than two groups were compared with Kruskal-Wallis test.

To evaluate the effect of the Cpt1a knock-in (CPT1A KI) strategy on renal damage we first used the FAN injury model (**Supplemental Figure 1C, upper panel**). While we found no significant differences in the degree of tubular atrophy and dilatation (**Supplemental Figure 2, A-C**), fibrosis was markedly ameliorated in Pax8-rtTA^tg/0^: tetO-Cpt1a^tg/0^ compared with WT mice (**Figure 2A, left panel**) and a significant reduction in the proportion of fibrosis as quantified by Sirius red was observed in the CPT1A KI model (**Figure 2A, right panel**). Renal function was reduced in FAN-treated WT animals, as reflected by the increase in blood urea nitrogen (BUN) and creatinine, but this effect was blunted in mice overexpressing CPT1A (**Figure 2, B and C**). The protein expression of classical profibrotic markers, fibronectin (FN) and smooth muscle actin (aSMA) was also significantly reduced in the CPT1A KI mice (**Figure 2, D and E**). To confirm the beneficial effect of FAO-gain-of-function in kidney fibrosis, the latter was assessed by performing UUO for 3 and 7 days in mice overexpressing CPT1A in TECs (**Supplemental Figure 1D**). Obstructed kidneys from WT animals showed significant tubulo-interstitial architectural and histological changes 7 days after UUO, while the extent of tubular atrophy and dilatation was markedly reduced in kidneys with increased levels of CPT1A (**Supplemental Figure 3A, left panel**). Evaluation of renal lesions by light microscopy showed increased collagen deposition in the interstitial area after 7 days of the procedure in the obstructed kidneys compared to the contralateral ones in WT mice. A significant protective effect of CPT1A overexpression was observed by a reduction of 20-40% in collagen deposition (**Supplemental Figure 3A, right panel**). As expected, circulating levels of BUN and creatinine were not different between WT and CPT1A KI mice after 7 days of UUO due to the remaining functional kidney (**Supplemental Figure 3, B and C**). A less dramatic but still significant decrease in FN and αSMA was also observed (**Supplemental Figure 3, D and E**).

**Figure 2.**
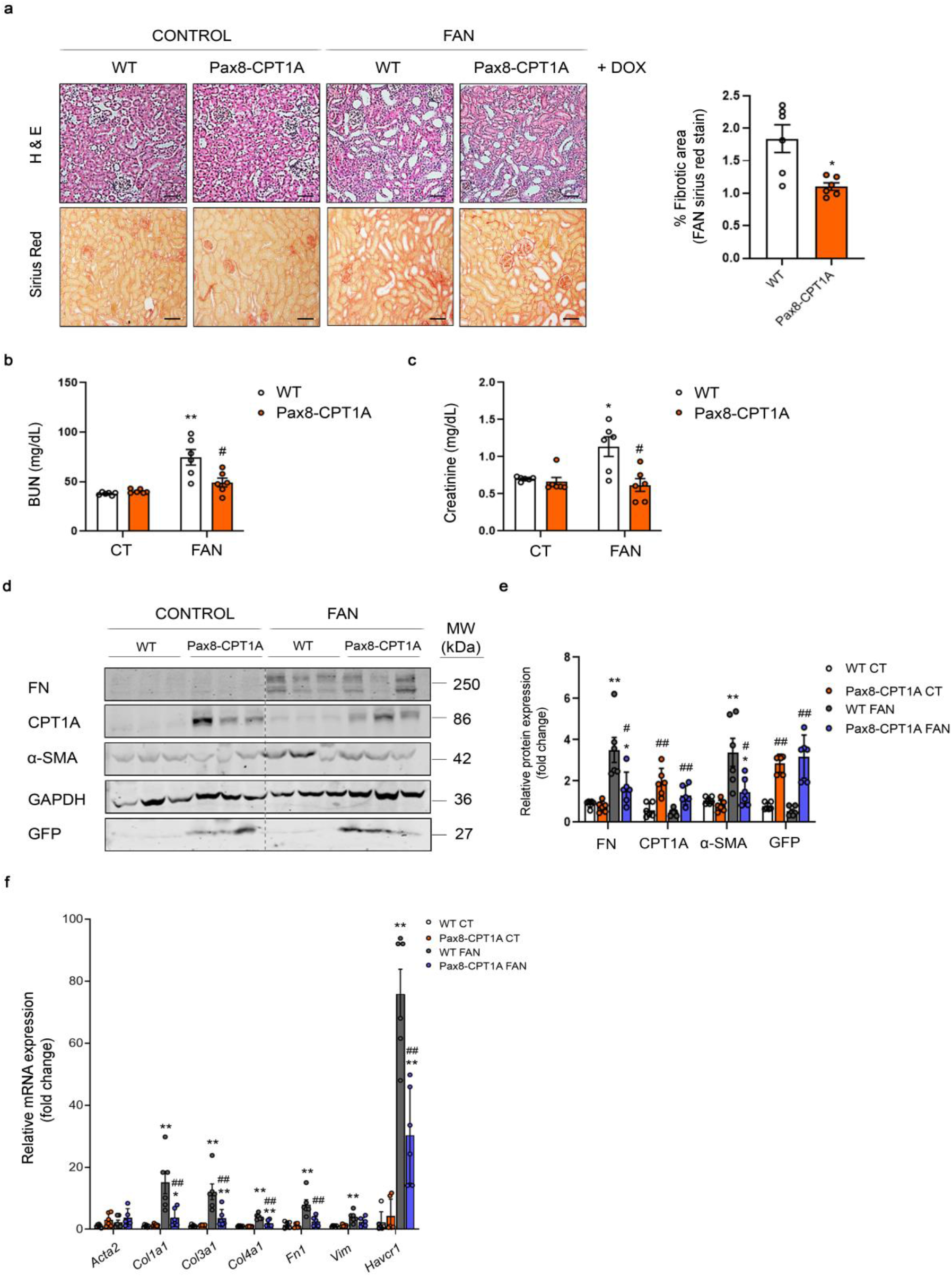
CPT1A overexpression prevents FAN-associated kidney function deterioration and experimental renal fibrosis. (**A**) Representative microphotographs from one mouse per group of hematoxylin and eosin (H&E) (upper panels) and Sirius Red (lower panels) staining of kidneys from WT and Pax8-CPT1A mice subjected to FAN after doxycycline treatment (Dox). Scale bars: 50 μm. Quantification of Sirius Red staining represents the mean ± s.e.m, n = 6 mice. *P < 0.05, compared to FAN/C kidneys in WT mice, respectively. (**B, C**) Serum blood urea nitrogen (BUN) (**B**) and serum creatinine (**C**) levels of WT and Pax8-CPT1A mice subjected to FAN after doxycycline treatment. Data represent the mean ± s.e.m (n = 6 mice). *P < 0.05, **P < 0.01 compared to its respective experimental control (CT) condition; #P < 0.05, compared to WT mice with the same experimental condition. (**D**) Immunoblots depicting fibronectin (FN), carnitine palmitoyltransferase 1A (CPT1A), green fluorescence protein (GFP) and alpha-smooth muscle actin (α-SMA) protein levels in kidneys from control (CT) and FA-treated (FAN) WT and Pax8-CPT1A mice after doxycycline induction. (**E**) Bar graphs represent the mean ± s.e.m. of fold changes corresponding to densitometric analyses (n = 6 mice). (**F**) mRNA levels of fibrosis-associated genes were determined by qRT-PCR using TaqMan qPCR probes in kidneys from control (CT) and FA-treated (FAN) WT and Pax8-CPT1A mice after doxycycline induction. Bar graphs represent the mean ± s.e.m. of fold changes (n = 6 mice). (E and F) *P < 0.05, **P < 0.01 compared to their corresponding control (CT) kidneys; ^#^P < 0.05, ^##^P < 0.01 compared to kidneys from WT mice with the same experimental condition. Statistical significance between two independent groups was determined using non-parametric two-tailed Mann-Whitney test, while more than two groups were compared with Kruskal-Wallis test. For gene nomenclature see **Supplemental Table 4**.

To understand the mechanisms underlying the protective action of CPT1A on renal fibrosis, we performed expression analysis of whole kidney from WT or CPT1A KI mice subjected to FAN or 7-days UUO. Expression of genes related to critical cellular functions for the initiation and perpetuation of tubular dysfunction and chronic tissue damage was analyzed by using specific TaqMan probes. A reduced expression for this subset of genes was observed in the FAN model. QRT-PCR-based quantification displayed lower expression of epithelial injury (*Havcr1-KIM-1*) and fibrotic markers (collagens) in kidneys from CPT1A KI mice (**Figure 2F**). These data were confirmed by mRNA quantitative analysis using Sybr green (**Supplemental Figure 4A**). A similar effect was observed in the UUO model (**Supplemental Figure 3F, and Supplemental Figure 4B**). These observations were also confirmed in the CKD mouse model of adenine-induced nephrotoxicity (ADN) (**Supplemental Figure 1E**). CPT1A KI mice showed a reduction in the expression of the crucial genes *Col1a1, Fn1* and *TGF-β* involved in these cellular mechanisms related to kidney fibrosis (**Supplemental Figure 5, A and B**) as well as the same trend in the injury marker KIM-1 (**Supplemental Figure 5, C and D**). Data collected from the three models of CKD in the CPT1A KI mice strongly support that CPT1A is an enzyme, which, by itself, has a crucial impact on the outcome of fibrosis most likely due to its critical function in the facilitation of FAO.

### CPT1A overexpression prevents mitochondrial dysfunction and restores FAO in the fibrotic kidney

Renal mitochondrial abnormalities and dysfunction are common features in the pathogenesis of different forms of renal disease (6). Cellular pathways promoting kidney damage can compromise mitochondrial homeostasis reflected in increased oxidative stress, apoptosis, microvascular loss and fibrosis, all of which contribute to renal function deterioration (7). Thus, we then evaluated morphological alterations of mitochondria in cortical proximal tubules by transmission electron microscopy. Cells from tubular segments of healthy kidneys presented regular apical microvilli, intact basement membrane and basal infoldings. Mitochondria were very abundant; most of them presented an elongated shape and were localized in the basolateral part of the cells. They displayed a well-defined arrangement of well-preserved mitochondrial cisternae with a homogeneous inner matrix. In contrast, in the FA-treated mice group, many epithelial cells were detached from the tubular basement membrane and showed disrupted basal infoldings. Mitochondrial structure was lost and mitochondria presented a fragmented, small and round appearance. Interestingly, most of these morphological alterations in mitochondria as well as the reduction in mitochondrial mass induced by FA were almost abrogated in renal epithelial cells overexpressing CPT1A (**Figure 3A**). CPT1A overexpression prevented the drop of mtDNA copy number in the 3-days UUO (**Supplemental Figure 6A**) and FAN (**Figure 3B**) models of kidney damage. As expected, defective FAO was observed in fibrotic kidneys from WT mice. However, in the 3-days UUO, FAN and ADN models, CPT1A overexpression counteracted this impairment, maintaining a FAO rate comparable to healthy kidneys (**Figure 3C, and Supplemental Figure 6, B and D**). Closely related to the improvement in FAO by CPT1A, ATP content in whole kidney tissue increased from 50 to 80 μM/mg protein after CPT1A overexpression. In the 3-days UUO, FAN and ADN models, CPT1A overexpression rescued the drop in ATP levels (**Fig.3D and Supplemental Figure 6, C and E**). These results suggest that appropriate levels of CPT1A and metabolic function are necessary and sufficient to preserve adequate mitochondrial architecture and morphology.

**Figure 3.**
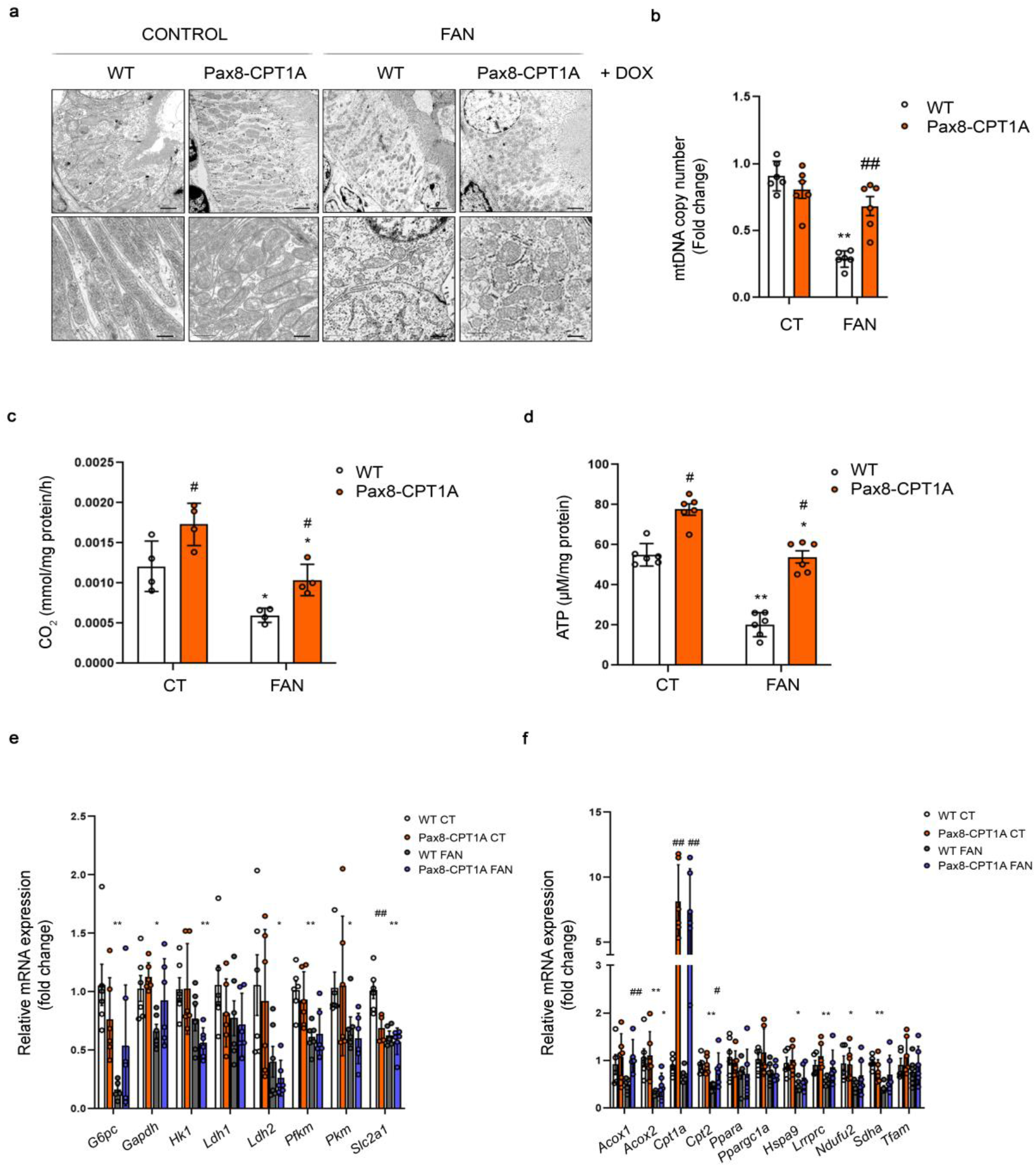
CPT1A prevents impaired mitochondrial morphology and fatty acid oxidation defect in FAN-induced kidney fibrosis. (**A**) Representative electron microscopy images of cortical proximal tubules from control and Pax8-CPT1A mice subjected to FAN after doxycycline induction. Scale bars: 10 μm (upper panels), 100 nm (lower panels). (**B**) Mitochondrial DNA copy number (mtDNA) was determined in kidneys of WT and Pax8-CPT1A mice in the FAN model. Bar graphs represent the mean ± s.e.m. of fold changes (n = 6 mice). **P < 0.01 compared to their corresponding control (CT) kidneys; ^##^P < 0.05 compared to kidneys from WT mice with the same experimental condition. (**C**) Radiolabeled palmitate-derived CO_2_ was determined after incubation of ^14^C-palmitate with kidney tissue from WT and Pax8-CPT1A mice in the FAN model after doxycycline induction. (**D**) ATP levels in total kidney tissue determined in mice subjected to FAN model. (C and D) Bar graphs represent the mean ± s.e.m (n = 4 mice). *P < 0.05, **P < 0.01 compared to their corresponding control (CT) kidneys; ^#^P < 0.05 compared to kidneys from WT mice with the same experimental condition. (**E, F**) mRNA levels of glucose utilization-(**E**) and peroxisomal/mitochondrial function-(**F**) associated genes were determined by qRT-PCR using TaqMan qPCR probes in kidneys from control (CT) and FA-treated (FAN) WT and Pax8-CPT1A mice after doxycycline induction. (E and F) Bar graphs represent the mean ± s.e.m. of fold changes (n = 6 mice). *P < 0.05, **P < 0.01 compared to their corresponding control (CT) kidneys; ^#^P < 0.05, ^##^P < 0.01, compared to kidneys from WT mice with the same experimental condition. Statistical significance between two independent groups was determined using non-parametric two-tailed Kruskal-Wallis test. For detailed gene nomenclature see **Supplemental Table 4**.

### CPT1A gain-of-function results in enhanced FAO-associated respiration of renal tubular epithelial cells even at the expense of glycolysis and AMPK activation

TECs use glucose for anaerobic glycolysis. Metabolic alterations of these cells during kidney fibrosis not only involve a defect in FAO but also in glucose oxidation (3). We found that in the FAN model there was a general downregulation trend in the expression of glycolysis-related genes, which was not recovered by CPT1A overexpression (**Figure 3E**). By contrast, FAN-induced repression in mRNA levels of the peroxisomal/mitochondrial function-related genes *Acox1, Cpt2, Lrpprc, Sdha* and *Tfam* was prevented in kidneys from CPT1A KI mice (**Figure 3F**). In the UUO model we found that levels of the majority of the analyzed regulators of glucose utilization were not altered in obstructed kidneys compared to contralateral ones. Only the increased expression of *Ldh1* and *Slc2a1* genes induced by UUO was prevented by CPT1A overexpression (**Supplemental Figure 6F**). Similarly, CPT1A-gain-of-function did not induce a major shift towards the expression of peroxisome/mitochondrial-related genes in contralateral kidneys (**Supplemental Figure 6G**).

To gain insight about quantitative metabolic changes at the cellular level we examined oxygen consumption (OCR) and extracellular acidification rates (ECAR) of primary TECs isolated from kidneys from CPT1A KI mice (**Figure 4A**). We found that basal and maximum OCRs were markedly higher when palmitic acid was supplied to TECs, indicating that TECs efficiently metabolize palmitate. The increase in OCR was sensitive to the CPT1 inhibitor Etomoxir, confirming its specificity (**Figure 4B**). FAO-associated OCR was also higher in primary kidney epithelial cells isolated from CPT1A KI compared to the ones isolated from WT mice. Cells treated with TGF-β1 had a lower baseline of oxygen consumption levels and showed a reduction in palmitate-induced elevation in OCR, indicating a low activity of fatty acid metabolism. CPT1A overexpression prevented the TGF-β1-induced bioenergetics derangement (**Figure 4C**). Simultaneously, glycolytic function associated to palmitate consumption was determined by ECAR. Oligomycin-induced blockage of OXPHOS, allows determining the maximum glycolytic capacity. We found that CPT1A overexpression promoted the inhibition of basal glycolytic function both in the presence and absence of TGF-β1 (**Figure 4D**). The increase in ATP levels related to CPT1A overexpression was also preserved under treatment with TGF-β1 (**Figure 4E**).

**Figure 4.**
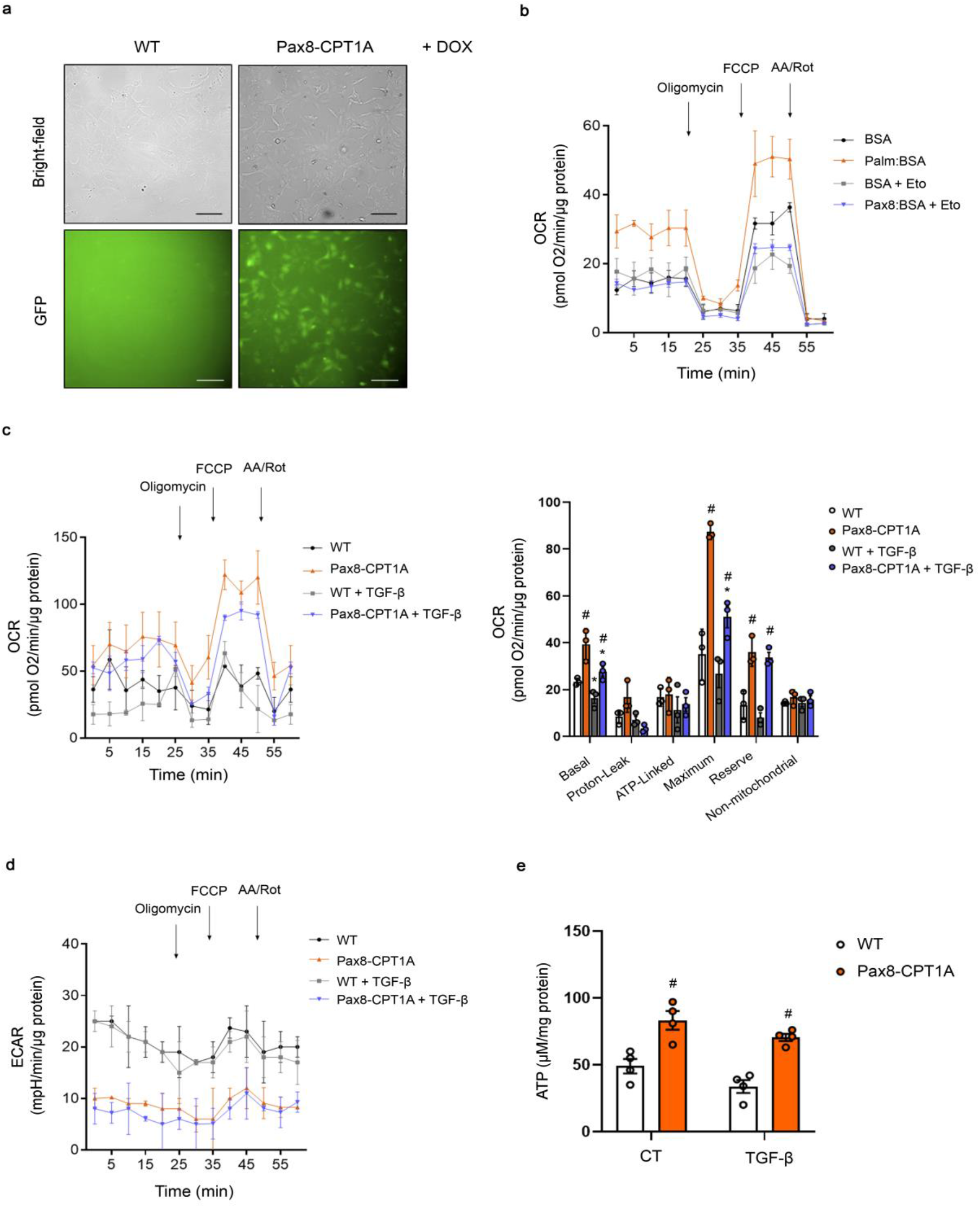
CPT1A overexpression prevents TGF-β1-induced FAO impairment and epithelial cell dedifferentiation. (**A**) Bright field or GFP immunofluorescence images of primary kidney epithelial cells (TECs) isolated from kidneys of WT and Pax8-CPT1A mice. Scale bar, 20 μm. (**B**) Oxygen consumption rate (OCR) of TECs from WT mice was measured with a Seahorse XF24 Extracellular Flux Analyzer. Where indicated, cells were pre-treated with palmitate-BSA FAO Substrate (200 μM) or the CPT1 inhibitor Etomoxir (Eto, 400 μM). Oligomycin (1 μM), FCCP (3 μM) and a combination of antimycin A (1 μM) and rotenone (1 μM) (AA/Rot) were injected sequentially at the time points indicated. Each data point represents the mean ± s.e.m of 4 independent experiments, each performed in triplicate. (**C**) Oxygen consumption rate (OCR) of TECs from WT and Pax8-CPT1A mice was measured with a Seahorse XF24 Extracellular Flux Analyser. Bar graphs (right panel) show the rates of OCR associated to basal, proton-leak, ATP-linked, maximum, reserve capacity and non-mitochondrial respiratory statuses. *P < 0.05 compared to their corresponding control (CT) TECs; ^#^P < 0.05 compared to TECs from WT mice with the same experimental condition. (**D**) Extracellular acidification rate (ECAR) of cells treated as in (A). Each data point represents the mean ± s.e.m of 4 independent experiments, each performed in triplicate. (**E**) ATP levels of TECs from WT and Pax8-CPT1A mice. Bar graphs represent the mean ± s.e.m (n = 4 mice per group). ^#^P < 0.05 compared to kidneys from WT mice with the same experimental condition. Statistical significance between two independent groups was determined using non-parametric two-tailed Kruskal-Wallis test.

To confirm the metabolic functional consequences of CPT1A overexpression in the human setting, we examined OCR and ECAR in the human tubular epithelial cell line, HKC-8. Adenoviruses carrying CPT1A (AdCPT1A) or adenoviruses control (AdControl) as a negative control were used to infect HKC-8 cells. CPT1A protein levels were 4-fold higher in CPT1A-expressing HKC-8 than in control cells (**Supplemental Figure 7, A and B**). In consistence with our observations in epithelial cells from murine kidneys overexpressing CPT1A, HKC-8 cells transduced with *CPT1A* exhibited a reduced TGF-β1-induced FAO inhibition (**Supplemental Figure 7C**) and suppression of glycolysis as reflected by ECAR levels (**Supplemental Figure 7D**). As expected, we found that the FAO rate was 2-fold higher in AdCPT1A-expressing HKC-8, as confirmed by measurement of ^14^C-palmitate-derived ^14^CO_2_ (**Supplemental Figure 7E**). Consistently, CPT1A overexpression increased the FRET signal of a specific ATP sensor and promoted a reduced decrement in the presence of TGF-β1 (**Supplemental Figure 7F**).

Depressed carbohydrate, amino acid and lipid oxidative pathways have been described in patients and animal models of CKD, leading to energy deprivation (6). AMPK is a highly conserved sensor of the intracellular metabolic status and plays a critical role in systemic energy homeostasis(8). To determine if FAO gain of function in kidneys of CPT1A KI mice was associated to changes in AMPK activation as a consequence of increased ATP levels, phosphorylation of AMPK was analyzed by immunoblot. In the FAN model, obstructed kidneys presented increased AMPK phosphorylation protein levels compared with contralateral kidneys. Importantly, increasing CPT1A levels attenuated AMPK phosphorylation in fibrotic kidneys in the FAN model (**Supplemental Figure 8, A and B**). The levels of phosphorylated acetyl-CoA carboxylase, directly dependent on AMPK, changed accordingly. Similar data were obtained in the UUO and adenine models (**Supplemental Figure 8, C-F**). Overall, we found that increased FAO associated to overexpression of CPT1A improves mitochondrial respiration in the context of reduced glycolysis. Moreover, it is likely that the enhancement of ATP production related to FAO reins in AMPK activation triggered by chronic kidney damage, with independence of the model employed.

### Overexpression of CPT1A modifies the cellular inflammatory profile and abrogates TGF-beta associated epithelial cell damage

Both studies in patients and in animal models show a strong correlation between infiltrated macrophage polarization and the extent of fibrosis (9). In the FAN model we found that overexpression of CPT1A correlated with a decreasing trend in macrophage influx (**Figure 5, A and B**) and flow cytometry analysis revealed a reduced proportion of renal pro-inflammatory M1 subpopulation (**Figure 5, C and D**). In the UUO (**Supplemental Figure 9, A and B**) and ADN (**Supplemental Figure 5, E and F**) models the CPT1A-induced decrement in macrophage influx was also confirmed. By contrast, the macrophage subpopulation positive for CD86 (M1) was higher than that for CD206 (M2) in fibrotic kidneys 3- and 7-days after UUO, while CPT1A overexpression did not affect this shift (**Supplemental Figure 9, C-F**). However, the M2 macrophage subpopulation was increased in fibrotic kidneys from CPT1A KI mice compared with the WT ones after 7 days UUO (**Supplemental Figure 9, E and F**). Infiltration of the macrophage subpopulation positive for both CD86 and CD206 (M1/M2) observed in obstructed kidneys from WT mice 3 or 7 days after UUO was also enhanced by CPT1A overexpression (**Supplemental Figure 9, D and F**). In addition, the abundance of this M1 subpopulation was lower in FAN-induced fibrotic kidneys than in UUO-associated ones (**Figure 5D, and Supplemental Figure 9, D and F**). These data suggest that the changes in the degree of macrophage infiltration and relative contributions of macrophage subpopulations are dependent on the model of kidney injury.

**Figure 5.**
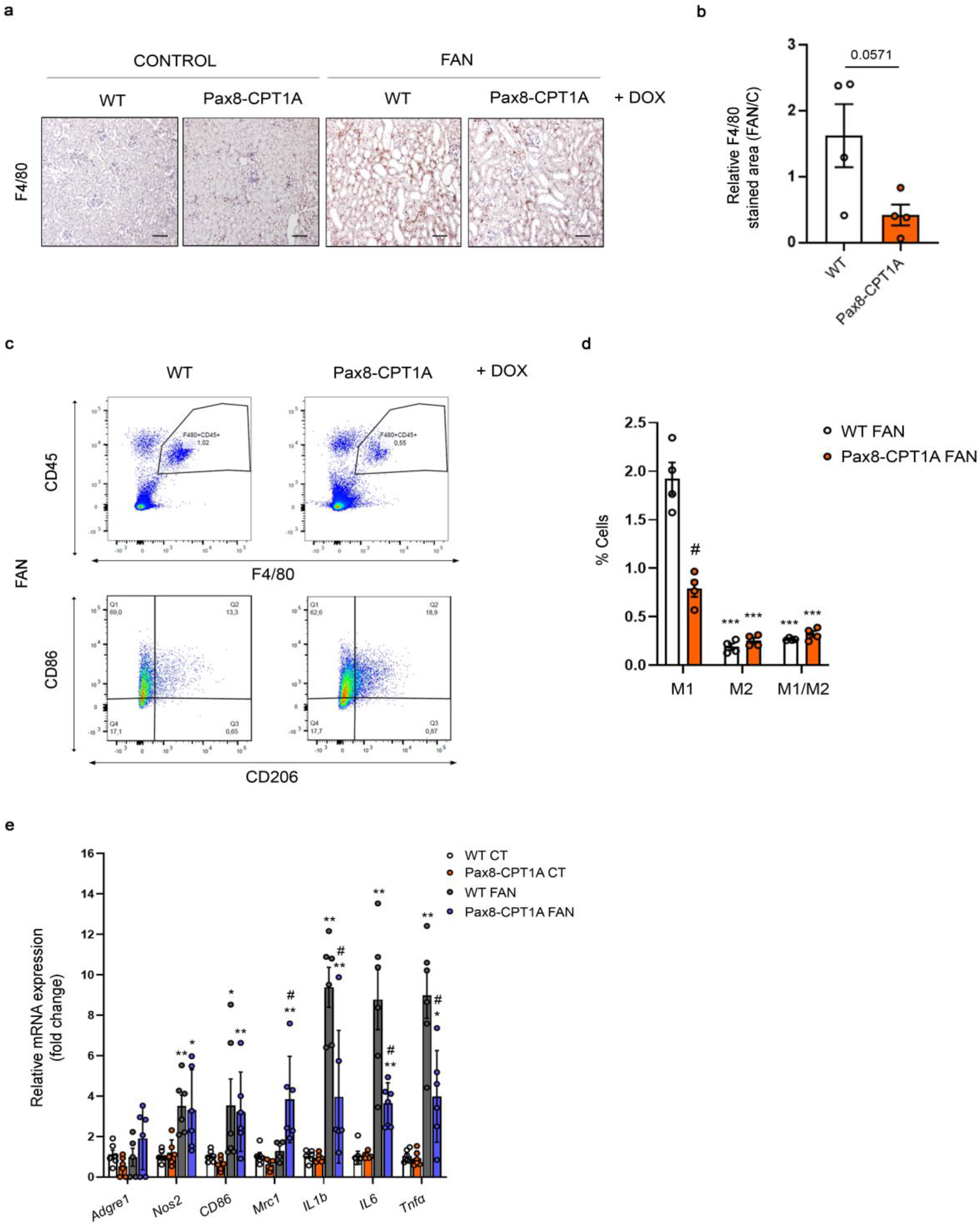
Overexpression of CPT1A reduces M1 macrophage infiltration in the FAN model. (**A**) Representative micrographs of one mouse per group showing the expression of F4/80 in kidney sections of mice treated as described above. Scale bar = 50 μm. (**B**) Bar graph represents the quantification of the % of F4/80 positive stained area in FA-treated mouse kidneys (FAN). Bar graphs represent the mean ± s.e.m, n = 4 mice. (**C**) Representative multiparameter flow cytometry dot plots showing the expression of CD45, and F4/80 in kidney cells from WT and Pax8-CPT1A mice subjected to FAN after doxycycline induction (upper panels) (one mouse per group is represented). CD86 and CD206 were used to determine the proportion of M1 and M2 macrophage subpopulations, respectively, in the total macrophage population (F4/80+, CD45+) (lower panels). Numbers in quadrants indicate cell proportions in percent of cells that express both markers. (**D**) Bar graph represents the percentage of kidney cells expressing CD86 (M1), CD206 (M2) or both (M1/M2) markers. Data represent mean ± s.e.m (n = 4 mice). ***P < 0.001 compared to M1 subpopulation in damaged kidneys from WT mice; ^#^P < 0.05 compared to corresponding cell subpopulation in damaged kidneys from WT mice. (**E**) mRNA levels of inflammation-associated genes were determined by qRT-PCR using TaqMan qPCR probes in kidneys from control (CT) and FA-treated (FAN) WT and Pax8-CPT1A mice after doxycycline induction. Bar graphs represent the mean ± s.e.m. of fold changes (n = 6 mice). *P < 0.05, **P < 0.01, compared to their corresponding control (CT) kidneys; ^#^P < 0.05 compared to kidneys from WT mice with the same experimental condition. Statistical significance between two independent groups was determined using non-parametric two-tailed Mann-Whitney test, while more than two groups were compared with Kruskal-Wallis test. For detailed gene nomenclature see **Supplemental Table 4**.

Inflammation is a major hallmark of renal fibrosis, especially in early stages (9). Consistently, kidney infiltration by CD3+ positive T lymphocytes after FAN was reduced in CPT1A KI compared with WT mice (**Supplemental Figure 2D, E**). Thus, we analyzed the profile of prototypical molecules related to inflammation in kidneys from mice overexpressing CPT1A in the context of renal damage. In the FAN model we found a lower expression of inflammation-related markers including the cytokines *IL1b, IL6* and *Tnfa* (**Figure 5E**), in contrast to findings in the 7 days UUO model (**Supplemental Figure 10C**). Consistently, cells from kidneys of mice overexpressing CPT1A presented a reduced population of damaged epithelial cells (CD45-EPCAM+ CD24+) (10) compared to those of WT mice in both the FAN and 7-days UUO models (**Figure 6, A and B, and Supplemental Figure 10, A and B**). In keeping, in the FAN model, the levels of the necroptosis markers Receptor-interacting serine/threonine-protein kinase (RIPK1), Mixed Lineage Kinase Domain Like Pseudokinase (MLKL) and phosphorylated RIPK-3 were significantly blunted in the presence of CPT1A overexpression (**Figure 6, C-E**). Moreover, there was also a reduced expression of the apoptotic markers *Apaf1, Bax and Bcl2* (**Figure 6F**). A similar pattern regarding the expression of apoptotic markers was observed in the 7 days UUO model (**Supplemental Figure 10D**).

**Figure 6.**
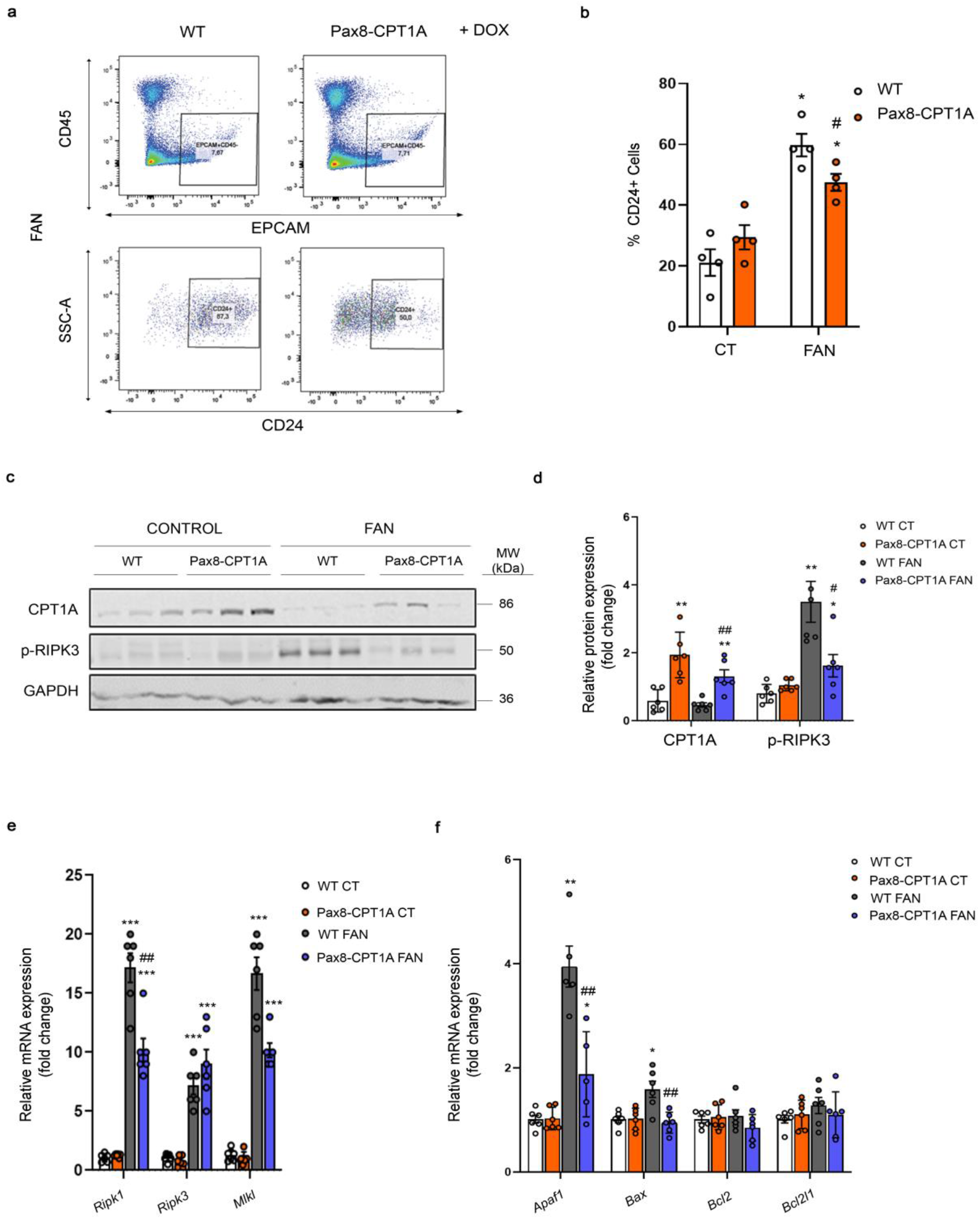
CPT1A overexpression reduces epithelial cell damage in the FAN model. (**A**) Representative flow cytometry dot plots from obstructed kidneys of WT and Pax8-CPT1A mice subjected to FAN after doxycycline treatment. Cells were gated for CD45 negative/Epithelial Cell Adhesion Molecule (EpCAM) positive (upper panels) and selected for the presence of CD24 (lower panels). Numbers in quadrants indicate cell proportions. (**B**) Bar graphs show the percentage of kidney cells positive for CD24. Data represent the mean ± s.e.m (n = 4 mice). *P < 0.05 compared to their corresponding control (CT) kidneys; ^#^P < 0.05 compared to damaged kidneys in WT mice. (**C, D**) Representative immunoblots and densitometries corresponding to CPT1A and phosphorylated RIPK3 protein levels in kidneys as in A (n=3 mice). *P < 0.05, **P < 0.01 compared to their corresponding control (CT) kidneys; ^#^P < 0.05, ^##^P < 0.01 compared to kidneys from WT mice with the same experimental condition. (**E**) mRNA levels of RIPK1, RIPK3 and MLKL were determined by qRT-PCR in kidneys as in A. Bar graphs represent the mean ± s.e.m. of fold changes (n = 6 mice). ***P < 0.001 compared to their corresponding control (CT) kidneys; ^##^P < 0.01, compared to kidneys from WT mice with the same experimental condition. (**F**) mRNA levels of apoptosis-associated genes were determined by qRT-PCR using TaqMan qPCR probes in kidneys from control (CT) and FA-treated (FAN) WT and Pax8-CPT1A mice after doxycycline induction. Bar graphs represent the mean ± s.e.m. of fold changes (n = 6 mice). *P < 0.05, **P < 0.01 compared to their corresponding control (CT) kidneys; ^##^P < 0.05 compared to kidneys from WT mice with the same experimental condition. Statistical significance between two independent groups was determined using non-parametric two-tailed Kruskal-Wallis test. For detailed gene nomenclature see **Supplemental Table 4**.

Epithelial cell dedifferentiation is a process associated to the transformation of a terminal cellular phenotype into one with relatively higher potential of differentiation into more than one cell type (11). This dedifferentiation is characterized by the loss of epithelial markers and acquisition of mesenchymal features, a process known as epithelial-to-mesenchymal transition (EMT) (12), the role of which in kidney fibrosis remains controversial. We studied the expression of E-cadherin, whose loss is considered as a representative feature of EMT, in the setting of CPT1A overexpression and found no significant differences in renal tubular expression in any of the three models employed (**Supplemental Figure 11A**). A similar pattern was observed when further EMT-associated markers were analyzed in the FAN model. Only the EMT-associated increased expression of Snail was prevented in the kidneys of CPT1A KI mice (**Supplemental Figure 11B**). We also tested whether enhancing renal epithelial FAO had a protective role in TGF-β-induced transformation of epithelial cells into a cellular fibrotic phenotype. Primary kidney epithelial cells isolated from CPT1A KI mice and treated with TGF-β1 for 48 h showed a marked reduction in the increase of these EMT-associated markers compared to the cells isolated from WT mice (**Supplemental Figure 11C**). Consistently, HKC-8 cells transduced with *CPT1A* exhibited a lower expression of collagen (**Supplemental Figure 11D**) and fibronectin (**Supplemental Figure 11E**). This set of data supports that the protective action of CPT1A in kidney fibrosis operates, at least in part, through the attenuation of TGF-β1-induced tubular epithelial cell transformation.

### Overexpression of CPT1A post damage markedly attenuates the fibrotic phenotype

To investigate if re-instauration of FAO-GOF after injury could also protect from kidney damage, we started doxycycline administration after FA injection and confirmed that CPT1A levels increased gradually in the CPT1A KI mice (**Supplemental figure 12, A - C**). We observed that the fibrotic area as well as the renal function impairment were significantly reduced in the CPT1A KI model (**Figure 7, A-C**). The expression of the profibrotic markers *Col1a1* and *Fn1* was also significantly reduced in the CPT1A KI mice (**Figure 7D**). TECs from CPT1A KI mice also exhibited a more favorable mitochondrial bioenergetics profile after damage, which was reflected by a higher ATP production rate and basal and maximum OCRs (**Figure 7, E and F**). This data reinforces the potential of FAO-GOF to combat kidney fibrosis even when renal damage has already been initiated.

**Figure 7.**
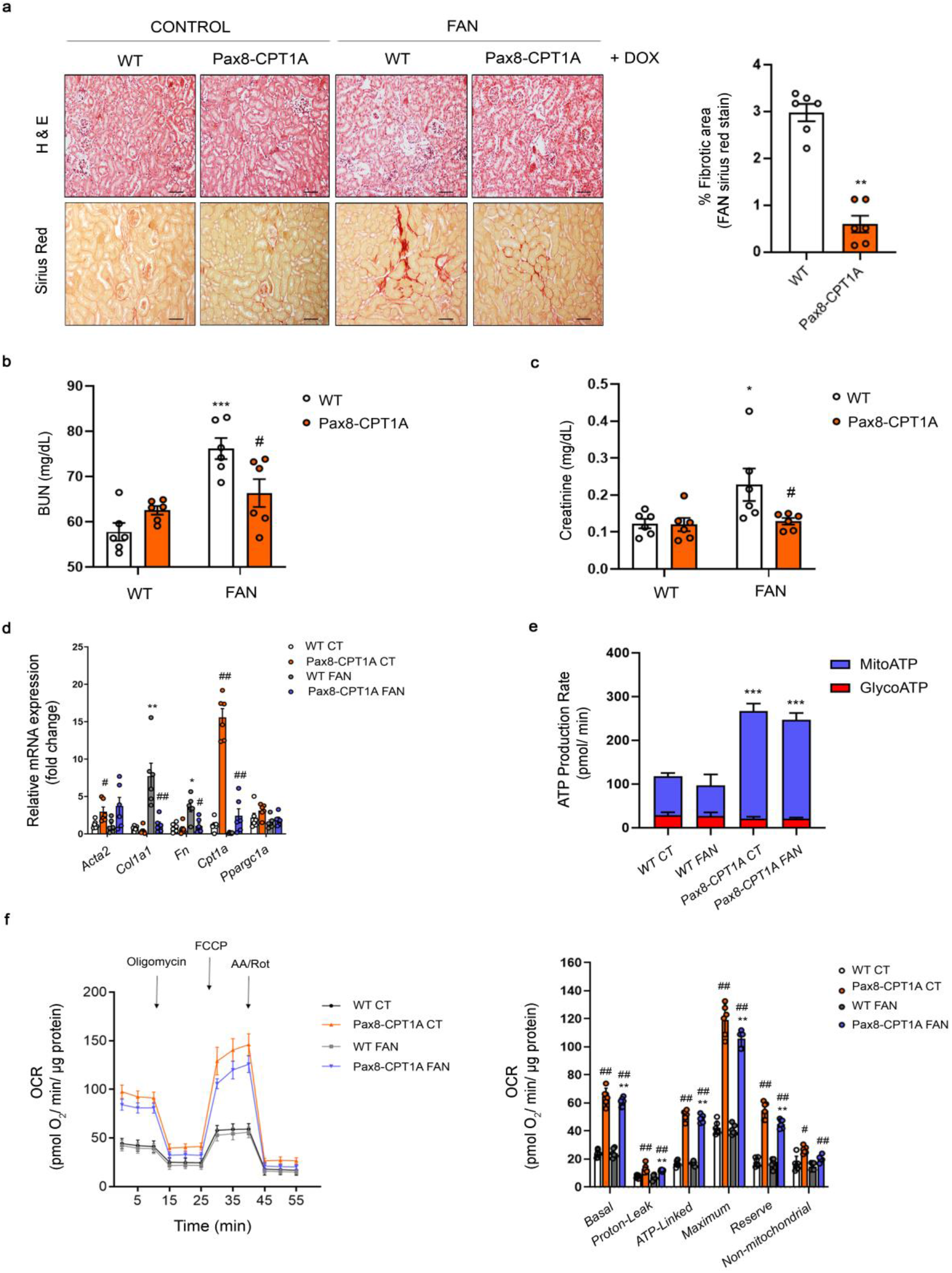
CPT1A upregulation after FA-induced renal disease mitigates FAN-associated kidney function deterioration, renal fibrosis and fatty acid oxidation defects. (**A**) Representative microphotographs of H&E (upper panels) and Sirius Red (lower panels) staining of kidneys from WT and Pax8-CPT1A mice subjected to FAN previous to doxycycline (Dox) (**Supplemental figure 12A**). Scale bars: 50 μm. Quantification of Sirius Red staining represents the mean ± s.e.m, n = 6 mice. **P < 0.05, compared to FAN kidneys in WT mice. (**B, C**) Serum BUN (**B**) and creatinine (**C**) levels of WT and Pax8-CPT1A mice subjected to FAN as in A. Data represent the mean ± s.e.m (n = 6 mice). *P < 0.05, ***P < 0.001 compared to its respective control (CT) condition; ^#^P < 0.05, compared to WT mice with the same condition. (**D**) mRNA levels of α-SMA, Col1α1, FN, CPT1A and Ppargc1a determined by qRT-PCR in kidneys of WT and Pax8-CPT1A mice subjected to FAN as in A. Bar graphs represent the mean ± s.e.m. of fold changes (n = 6 mice). *P < 0.05, **P < 0.01 compared to their corresponding control (CT) kidneys; ^#^P < 0.05, ^##^P < 0.01, compared to kidneys from WT mice with the same condition. (**E, F**) ATP production rate of TECs from WT and Pax8-CPT1A mice subjected to FAN as in A. (**F**) Oxygen consumption rate (OCR) of TECs from WT and Pax8-CPT1A mice subjected to FAN as in A. Bar graphs (right panel) show the rates of OCR as in Figure 4C. *P < 0.05 compared to their corresponding control (CT) TECs; ^#^P < 0.05 and ^##^P < 0.01 compared to TECs from WT mice with the same condition. Statistical significance between two independent groups was determined using non-parametric two-tailed Mann-Whitney test, while more than two groups were compared with Kruskal-Wallis test.

### An increase in plasma acyl-carnitines and a reduction of CPT1A levels is present in patients with CKD

To explore the importance of FAO in a clinical context we analyzed a cohort of 686 patients with CKD and diabetes pertaining to the PREDIMED study (13) (see **Supplemental Table 7** for details). In those where renal function parameters were available (n=686), we determined the levels of acyl-carnitines. We found inverse correlation between GFR and short (c2-c7) and medium (c8-c14)-chain acyl-carnitine levels (**Fig. 8A, 8B and Supplemental Table 8 and 9**) in patients with a GFR under 60 ml/min. Thus, CKD patients with a higher GFR showed less accumulation of short and medium-chain acyl-carnitines (**Fig. 8D, E, G**) with no correlation in the case of long-chain (c16-c26) acyl-carnitines (**Fig. 8C, F**). In a different cohort of CKD patients (see **Supplemental Table 10** for details) we found a positive correlation between tubule CPT1A expression levels and eGFR (**Figure 8, H and I**) in a RNAseq study from different pathological backgrounds. The degree of fibrosis also correlated significantly with declining CPT1A levels (**Figure 8J**). The increase in the levels of acyl-carnitines and the reduction in the levels of the limiting step enzyme responsible for their metabolism (CPT1A) most likely reflect a decreased FAO capacity associated to CKD. Thus, these results underscore the relevance of reduced FAO in CKD and support the importance of its gain-of-function to combat kidney fibrosis as demonstrated in our experimental model.

**Figure 8.**
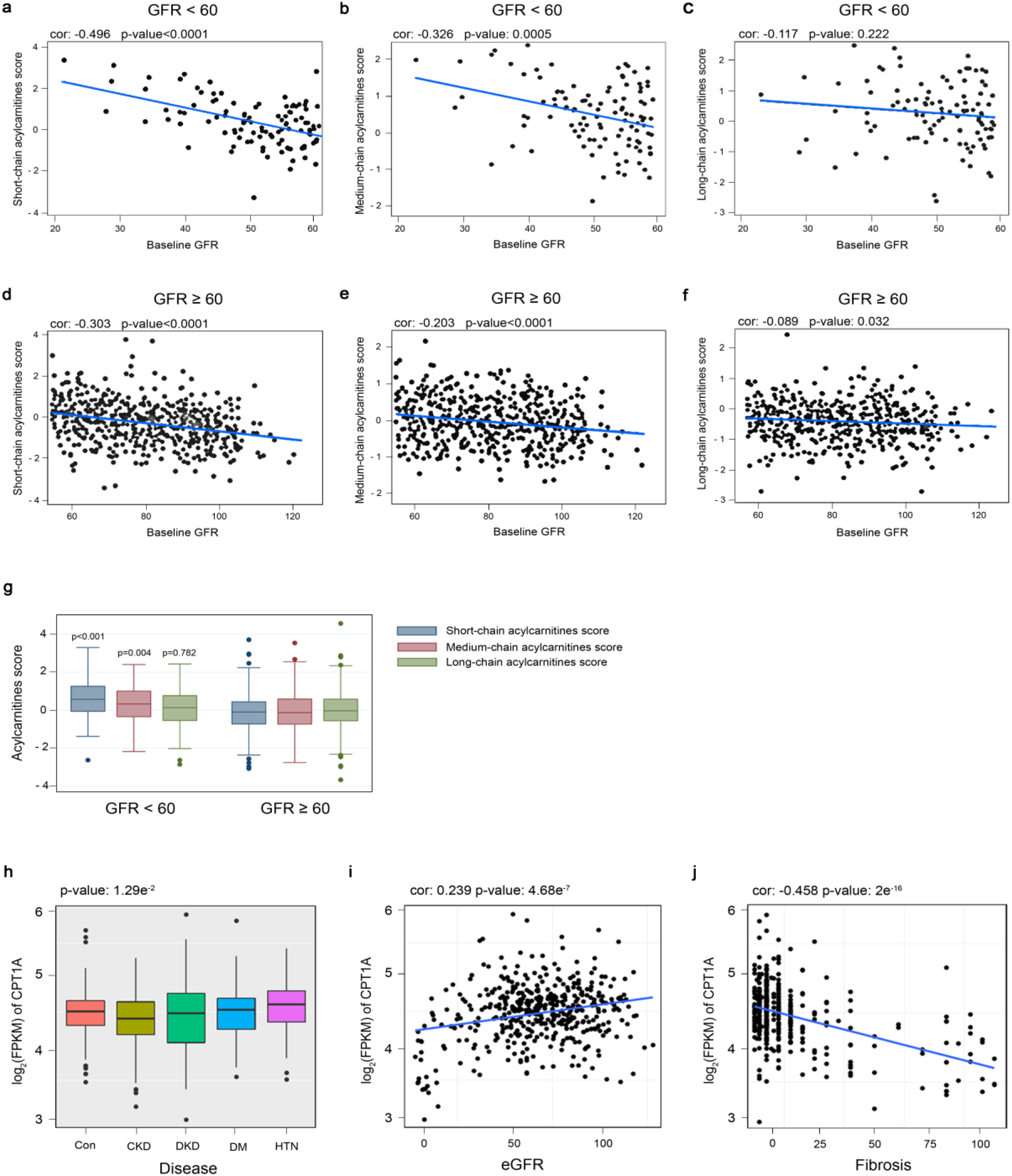
Plasma acyl-carnitines and CPT1A levels in patients with CKD. (**A-C**) Correlation between baseline GFR values and plasma short-chain acyl-carnitines (**A**), medium-chain acyl-carnitines (**B**) and long-chain acyl-carnitines (**C**) score in CKD patients with GFR<60. (**D-F**) Correlation between baseline GFR values and plasma short-chain acyl-carnitines (**D**), medium-chain acyl-carnitines (**E**) and long-chain acyl-carnitines (**F**) score in CKD patients with GFR≥60. (**G**) Baseline acyl-carnitine levels by CKD stage. P values for the comparison of acyl-carnitine levels between participants with GFR>60 vs GFR≥60. (**H**) CPT1A levels in renal biopsies from control (con) and patients with chronic kidney disease (CKD), diabetic kidney disease (DKD), diabetes mellitus (DM) or hypertension (HTN). (**I, J**) Correlation between CPT1A kidney levels and eGFR (**I**) or fibrosis score (**J**). For box-and-whiskers plots (G, H), within each box, horizontal white lines denote median values; boxes extend from the 25^th^ to the 75^th^ percentile of each group’s distribution of values; vertical extending lines denote adjacent values (i.e., the most extreme values within 1.5 interquartile range of the 25^th^ and 75^th^ percentile of each group); dots denote observations outside the range of adjacent values. (A - G) Chi-squared and Student’s t-test were used to compare categorical and quantitative variables, respectively. (H) ANOVA test was used to assess the significance across different disease groups. Cor.test() function in R was used to get the Pearson’s correlation and the corresponding P-values.

## Discussion

Kidney fibrosis is critically linked to metabolic failure in tubular epithelial cells (14). We demonstrate that genetic gain-of-function of FAO is sufficient to provide significant mitigation of fibrosis development in three different experimental models of chronic kidney damage. Moreover, FAO-GOF is also therapeutically effective when reinstated after kidney damage. Although FAO is a complex process depending on several biochemical steps and enzymatic systems, CPT1A is rate-limiting due to its key role for shuttling medium and long acyl CoA chains into the mitochondrial matrix with the concourse of L-carnitine (15). To date, no drugs are available to specifically activate CPT1 and hence, pharmacological studies are not devoid of limitations. Homozygous CPT1A deficiency is lethal in the mouse (16), while tissue specific CPT1A knockout models for the endothelium (17), pancreatic α cells (18) and intestinal stem cells(19) have been generated. None of these studies addressed potential consequences on organ fibrosis. The FAO gain-of-function genetic model herein described is likely one of the first direct demonstrations of the potential of this strategy to treat kidney fibrosis. The study was performed in heterozygous mice in an attempt to avoid non-physiological scenarios due to CPT1A overdosing (20). Mitigation of renal fibrogenesis was reflected in a reduction of fibrotic markers, histological improvement and amelioration of renal function. Slight differences in the magnitude of changes are most likely due to the variation in the intensity of the inflammatory or the fibrotic components among models. While no experimental model recapitulates with close fidelity human CKD, the significant correlation between advanced kidney disease and two signatures of FAO reduction (increased acyl-carnitines and low CPT1A) in two large patient cohorts attests to the clinical relevance of a FAO-related major metabolic disturbance in human CKD (21). Adjustment for age, sex, diabetic state and albumin/creatinine ratio supports that renal function was likely the variable accounting for these differences, even though the contribution of other tissues, such as adipose, cannot be completely excluded. The fact that we could not evaluate levels of CPT1A and acyl-carnitines simultaneously in any of the two cohorts poses some limitations for a clear-cut interpretation. We found correlation between GFR and short and medium but not long-chain acyl-carnitines levels, as could be initially expected. In a detailed study, Afshinnia et al. found higher plasma abundance of long chain free FAs coupled with a significantly lower long-to-intermediate acyl-carnitine ratio (a marker of impaired *β*-oxidation) in severe stages of CKD, but CPT1A levels were not determined (21). Increased levels of short and middle ACs in the diabetic cohort with CKD may be explained by alterations in the expression or activity of FAO enzymes aside CPT1A(3) as short and middle ACs may derive from long chain ACs incomplete fatty acid oxidation (22). Further, the systemic clearance of ACs may be differentially impaired in this population, also impacting on their biodistribution (23, 24).

The high energetic demand of renal tubular epithelial cells, required for their performance of multiple functions related to the homeostasis of the internal milieu, dictates their dependence on an intact mitochondrial function. We show that FAO gain-of-function, attained by overexpressing tubular CPT1A (in both animals and cells) restores mitochondrial mass and architecture, enhances OCR and increases ATP production in conditions of renal damage. This appears to occur at the expense of glycolysis as ECAR is significantly lower under conditions of CPT1A overexpression. The reciprocal regulation between FAO and glycolysis, the Randle cycle (25), has been demonstrated in muscle and heart, our data now supporting that it also takes place in renal tubular cells. Alternatively, it is possible that TECs are consuming lactate as the main fuel for gluconeogenesis, a process in which the kidney plays an essential role (26). The fundamental question as to whether metabolism dictates phenotype is in part answered here, as our data support that an increased capacity of the tubular epithelial cell to meet its energy challenges is able to both prevent and revert kidney fibrosis. Thus, efforts directed at enhancing FAO in the early stages of fibrosis should very likely pay off to combat human CKD.

## Materials and Methods

### Generation of a transgenic mouse model for inducible Cpt1a gene

Generation of an inducible conditional transgenic mouse model for *Cpt1a* overexpression in the renal epithelium was based on a 2nd generation Tet-On system with site-specific recombination in embryonic stem (ES) cells (27, 28). Mice harboring the transgene of *Cpt1a* gene under the control of the tetracycline-responsive promoter element (TRE) (tetO-Cpt1a mice) were generated in the transgenic mice core unit from the National Cancer Research Center (CNIO, Spain). Mice were generated by diploid blastocyst injection. Blastocysts were harvested at 3.5 days post coitum from C57/BL6-J strain females. Between 10 and 15 KH2 ES cells were injected per blastocyst. Approximately 30 injected blastocysts were transferred into pseudo-pregnant CD1 strain recipient females (transgenic mice core unit, CNIO, Spain). Chimeras from the litter with a high percentage agouti coat colour (>80%) were then crossed with C57/BL6-J mice to evaluate germline transmission. Next, these mice were crossed with mice providing renal epithelial tissue specificity, Pax8-rtTA (Jackson labs, Bar Harbor, Ma). Male tetO-Cpt1a^tg/0^ mice were bred with Pax8-rtTA^tg/0^ female mice to generate double heterozygous Pax8-rtTA^tg/0:^tetO-Cpt1a^tg/0^ mice and their control littermates in a B6N;129Sv mixed genetic background. Extra-renal expression only occurs at the level of the thyroid and this fact does not interfere with any kidney phenotype (5).

### Molecular cloning and gene targeting in ES cells

The CPT1A transgene containing the 5’UTR of the *Cpt1a* gene, the Cpt1a open reading frame (ORF) (Genbank accession number NM_013495.2), the 2A self-cleaving peptide (P2A), the gene encoding for green fluorescent protein (GFP) and the 3’UTR of the Cpt1a gene were cloned into a unique EcoRI site of pBS31 vector (OriGene Technologies, Rockville, Maryland). This vector contains the phosphoglycerate kinase (PGK) promoter followed by an ATG start codon and a FRT recombination site, a splice acceptor-double polyA cassette, the tetracycline operator with a cytomegalovirus (CMV) minimal promoter and an SV40 polyA signal, assembled in Bluescript. By using FRT/flippase-mediated recombination system(27, 28), the CPT1A transgene was targeted into the downstream region of the collagen 1a1 (Col1a1) locus of KH2 embryonic stem (ES) cells containing a frt-flanked PGK-neomycin-resistance gene followed by a promoterless, ATG-less hygromycin-resistance gene. These cells also contained the M2-rtTA under control of the endogenous ROSA26 promoter(29). For gene targeting, 50 μg pBS31 vector and 25 μg vector encoding the flippase (pCAGGS-FLP) (30) were co-electroporated with 1.5 x 107 KH2 ES cells at 400 V and 125 μf using two pulses in a Gene PulserII (Bio-Rad, Hercules, CA). Recombination between FRT sites such that the entire plasmid is inserted within the PGK promoter and the ATG initiation codon upstream and in frame with the hygromycin resistance gene was used to select correctly targeted cells. ES cells were treated with 140 μg/ml hygromycin B (Carl Roth, Karlsruhe, Germany) after 48 h post electroporation to select clones that had undergone site-specific recombination and individual clones were picked after 8-14 days(31). Individual ES clones were tested in vitro by treating them with 1μg/ml doxycycline (Sigma, St. Louis, MO) in the culture media for 4 days. Then, GFP was measured in a BD FacsCantoTM II cytometer (BD Bioscience, San José, CA) to assess the electroporation efficiency and cells expressing GFP were sorted.

### Doxycycline induction

To induce CPT1A expression, 8-week-old Pax8-rtTA^tg/0^:tetO-Cpt1a^tg/0^ mice and corresponding Pax8-rtTA^0/0^:tetO-Cpt1a^tg/0^ (WT) mice were fed with doxycycline (Sigma, St. Louis, MO) at concentrations of 1 mg/ml via drinking water for 3 weeks. Mice were housed in colony cages with a 12:12-h light-dark cycle in a temperature- and humidity-controlled environment with free access to water.

### Genotyping

Mice were genotyped by PCR using DNA extracted from tail biopsies and the primers listed in **Supplemental Table 1**. DNA extraction was performed by using the AccuStart™ II Mouse Genotyping Kit according to the manufacturer’s instructions.

### Cell lines and culture conditions

Human proximal tubular epithelial cells (HKC-8) were cultured in DMEM/F12 (Dulbecco’s modified Eagle’s medium 1:1 (v/v)) (Corning, New York, NY) supplemented with 15 mM Hepes, 5% (vol/vol) fetal bovine serum (FBS) (HyClone Laboratories, Logan, UT), 1x Insulin-Transferrin-Selenium (ITS) (Gibco, Rockville, MD), 0.5 μg/ml hydrocortisone (Sigma, St. Louis, MO), 50 units/mL penicillin and 50 μg/mL streptomycin (Gibco, Rockville, MD). This cell line was kindly provided by Dr. Susztak’s lab (Philadelphia, Pennsylvania, USA). HEK293A cells obtained from ATCC (Manassas, VA, USA) were cultured in DMEM supplemented with 10% (v/v) FBS (HyClone Laboratories, Logan, UT) and 1% penicillin /streptomycin (Gibco, Rockville, MD). All cells were cultured at 37°C and 5% CO_2_ and treated with trypsin every five days for sub-culturing. Treatments with human recombinant 10 ng/ml TGF-β1 (R&D Systems, Minneapolis, MN) were performed after serum-free starvation for 12 hours (h).

### Isolation of primary kidney epithelial cells

Kidneys from CPT1A KI and wild-type (WT) mice (3- to 5-week-old males) were collected after sacrifice minced into pieces of approximately 1 mm^3^. These pieces were digested with 10 ml HBSS containing 2 mg/mL collagenase I (Thermo Scientific, Rockford, IL) for 30 minutes at 37°C with gentle stirring and supernatants were sieved through a 100-μm nylon mesh. After centrifugation for 10 minutes at 3000 rpm, the pellet was resuspended in sterile red blood cell lysis buffer (8.26 g NH4Cl, 1 g KHCO3, and 0.037 g EDTA per 1 L ddH2O) and seeded in 10-cm culture dishes. Cells were cultured in RPMI 1640 (Corning, New York, NY) supplemented with 10% FBS (HyClone Laboratories, Logan, UT), 20 ng/mL EGF (Sigma, St. Louis, MO), 20 ng/ml bFGF (Sigma, St. Louis, MO), 50 units/mL penicillin and 50 μg/mL streptomycin (Gibco, Rockville, MD) at 37°C and 5% CO_2_. Cells were used between days 7 and 10 of culture.

### Immunoblot

Cells or a quarter piece of each kidney sample were homogenized and lysed in 100/300 μL RIPA lysis buffer containing 150 mM NaCl, 0.1% SDS, 1% sodium deoxycholate, 1% NP-40 and 25 mM Tris–HCl pH 7.6, in the presence of protease (Complete, Roche Diagnostics, Mannheim, Germany) and phosphatase inhibitors (Sigma-Aldrich, St. Louis, MO). Samples were clarified by centrifugation at 10,000 g for 15 min at 4°C. Protein concentrations were determined by the BCA Protein Assay Kit (Thermo Scientific, Rockford, IL) and was measured in Glomax^®^-Multi Detection system (Promega, Madison, WI). Equal amounts of protein (10–50 μg) from the total extract were separated on 8–10% SDS–polyacrylamide gels and transferred onto nitrocellulose blotting membranes (GE Healthcare, Chicago, IL) at 12 V for 20 min in a semi-dry Trans-Blot Turbo system (Bio-Rad, Hercules, California). Membranes were blocked by incubation for 1 h with 5% non-fat milk in PBS containing 0.5% Tween-20 and blotted overnight with the specific antibodies listed in **Supplemental Table 2**. After incubation with IRDye 800 goat anti-rabbit and IRDye 600 goat anti-mouse (1:15,000, LI-COR Biosciences, Lincoln, NE) secondary antibodies, membranes were imaged with the Odyssey Infrared Imaging System (LI-COR Biosciences, Lincoln, NE). Band densitometry was performed using the ImageJ 1.48 software (http://rsb.info.nih.gov/ij) and relative protein expression was determined by normalizing to GAPDH. Fold changes were normalized to values of control condition.

### RNA extraction

Total RNA was extracted from HKC-8 or mouse kidneys using the miRNeasy Mini Kit (Qiagen, Valencia, CA) according to the manufacturer’s instructions. RNA quantity and quality were determined at 260 nm by a Nanodrop-1000 spectrophotometer (Thermo Scientific, Rockford, IL).

### Analysis of mRNA expression

Reverse transcription (RT) was carried out with 500 ng of total RNA using the iScript™ cDNA Synthesis kit (Bio-Rad, Hercules, CA). qRT–PCR was carried out with the iQ™SYBR Green Supermix (Bio-Rad, Hercules, CA), using a 96-well Bio-Rad CFX96 RT–PCR System with a C1000 Thermal Cycler (Bio-Rad, Hercules, CA) according to the manufacturers’ instructions. A C_t_ value was obtained from each amplification curve using CFX96 analysis software provided by the manufacturer. Relative mRNA expression was determined using the 2^−ΔΔCt^ method (32). The *18S* gene was used for normalization purposes. The primer sequences used for mRNA quantification are listed in **Supplemental Table 3**. Fold changes were normalized to values of control condition.

### Mouse models of kidney fibrosis

Mice were housed in the Specific-pathogen-free (SPF) animal facility at the CBMSO in accordance with EU regulations for all the procedures. Unilateral ureteral obstruction, folic acid nephropathy and adenine induced nephrotoxicity models are described in **Supplemental Methods**.

### [1-14C] Palmitate oxidation

Measurement of fatty acid oxidation (FAO) rates was performed in mouse kidney tissue and cells as previously described (33). Kidneys from renal fibrosis mouse models were homogenized in 5 volumes of chilled STE buffer by a Dounce homogenizer. Thirty μL of tissue homogenate supernatant were mixed with 370 μL of the oxidation reaction mixture containing 7% BSA/5 mM palmitate/0.01 μCi/μL ^14^C-palmitate (PerkinElmer, Waltham, MA) and incubated at 37 °C for 30 min. In the case of cells, they were seeded in 12-well dishes to reach a confluence of 70% and were infected with CPT1A adenoviruses as described in the adenovirus-mediated CPT1A overexpression procedure section. Then, cells were incubated in 500 μL of media containing 0.3% BSA/100 μM palmitate/0.4 μCi/mL ^14^C-palmitate at 37 °C for 3 h. Each sample was assayed in triplicate. The reaction was stopped by the addition of 200 μl of 1 M perchloric acid. The rate of palmitate oxidation was measured as released ^14^CO_2_ trapped in a filter paper disk with 20 μL of 1 M NaOH in the top of sealed vials. The remaining acid solution for each sample was centrifuged at 14,000 g for 10 min at 4 °C and 400 μL of supernatant was also added to scintillation vials for the quantification of ^14^C-palmitate-derived acid-soluble metabolites. ^14^C products were counted in an LS6500 liquid scintillation counter (Beckman Instruments Inc., Brea, CA). Scintillation values were converted to mmol ^14^CO_2_ or acid-soluble metabolites multiplying the specific activity and normalized to the protein content.

### TaqMan gene expression assay

TaqMan Array Plates (Thermo Scientific, Rockford, IL) were selected as the platform for gene expression profiling. This panel consisted of a total of 43 unique TaqMan probes specific for mouse gene related with fibrosis, apoptosis, mitochondrial metabolism, glucose utilization and inflammation listed in **Supplemental Table 4**. RNA from kidney samples was performed as described in the RNA extraction procedure section. Two μg of total RNA were subjected to reverse transcription using High-Capacity cDNA Reverse Transcription Kit (Thermo Scientific, Rockford, IL) according to the manufacturer’s instructions. PCR amplification was performed using TaqMan^®^ master (Thermo Scientific, Rockford, IL) with the Roche LightCycler 480 Real-Time PCR system (AB7900HT). This procedure was performed in the Genomic Facility of the Fundación Parque Científico de Madrid (Madrid, Spain). The ABI TaqMan SDS v2.4 software was utilized to obtain Cq values (Cq) for each gene. The Cq data were analyzed with the StatMiner 4.2.8 software (Integromics; Madrid, Spain). The delta C_t_ (ΔC_t_) value was calculated by normalizing C_t_ values to the endogenous housekeeping 18S. Relative mRNA expression was determined using the 2^−ΔΔCt^ method (32). Fold changes were normalized to values of control condition.

### Mitochondrial copy number determination

Genomic DNA was extracted from mouse kidneys using the DNeasy Blood & Tissue Kit (Qiagen, Valencia, CA) according to the manufacturer’s instructions. Mitochondrial abundance was determined with the Mouse Mitochondrial DNA Copy Number Assay Kit (Detroit R&D, Detroit, MI). Relative mtDNA copy number was presented as the mtDNA-to-nuclear DNA ratio.

### Assessment of kidney function

Animal kidney function was determined by analyzing the indicators, serum creatinine and Blood Urea Nitrogen (BUN). They were analyzing by using the QuantiChrom TM Creatinine Assay Kit (Bioassay systems, Hayward, CA) and the Blood Urea Nitrogen (BUN) Colorimetric Detection Kit (Ann Arbor, MI), respectively, according to the manufacturers’ instructions. In the case of the reversion experiments (Fig.7), serum creatinine was measured by capillary electrophoresis (Agilent 7100) coupled to time of flight mass spectrometry (6224 Agilent) (CE-TOF-MS) (34, 35). A calibration curve was used for quantification with creatinine including methionine sulphone as internal standard.

### Measurement of ATP level

ATP content in kidney tissue, HKC-8 and primary kidney epithelial cells was measured by using the ATP colorimetric/fluorometric assay Kit (Biovision Inc, Milpitas, USA) according to the manufacturer’s instructions. Data were normalized for total protein content. In HKC-8 cells, relative ATP content was also assessed with the FRET Clover-mApple ATP sensor (36, 37). Cells were seeded in 12-well dishes to reach a confluence of 70% and transfected with 1.5 μg vector/well using lipofectamine 2000 (Invitrogen, Carlsbad, CA). Then, cells were infected with CPT1A adenoviruses as described in the adenovirus-mediated CPT1A overexpression procedure section and FRET signal was analyzed by flow cytometry. Cells were gated by forward (FSC-A) and side scatter (SSC-A), then for single cells using FSC-A/FSC-H. Next, dead cells were excluded. Fluorescence intensity was measured by flow cytometry using wavelengths (ex/em) 550/610 (acceptor), 490/550 (donor), and 490/600 (FRET) in a BD FacsCantoTM II system (BD Bioscience, San José, CA) and analyzed with the FlowJo 10.2 software (FlowJo, LLC, Ashland, OR). The ratio of FRET/Donor was displayed as a histogram and its median of fluorescence intensity was obtained. For each experimental condition, at least 20,000 singlets were analyzed in triplicates.

### Quantification of kidney cell populations by flow cytometry

Multiparametric flow cytometry was used for the identification of macrophage and epithelial cell population. Kidneys were diced, incubated at 37°C for 30 min with 0.5 mg/ml liberase DL (Roche Basel, Switzerland) and 100 U/ml DNase (Roche Basel, Switzerland) in serum free DMEM and filtered (40μm). For staining, 10^6^ cells were dissolved in 50 μl of buffer and pre-incubated with 0.25μl CD16/CD32 (BioLegend, San Diego, CA). Three μl of 4,6-diamidino-2-phenylindole (DAPI) (1:15000) (Sigma, St. Louis, MO) was added to stain dead cells. Cell suspensions were incubated with specific fluorochrome-conjugated antibodies (BioLegend, San Diego, CA) listed in **Supplemental Table 5**. For each experiment, flow minus one (FMO) controls was performed for each fluorophore to establish gates by using corresponding antibodies listed in **Supplemental Table 5**. Fluorescence intensity was measured in a BD FacsCantoTM II cytometer (BD Bioscience, San José, CA) and analyzed with the FlowJo 10.2 software (FlowJo, LLC, Ashland, OR). For each kidney sample, at least 20,000 singlets were analyzed in triplicates. Cell gating was initially based in the Forward versus Side scatter (FSC vs SSC) plot. Next, dead cells were excluded. Identification of the macrophage cell population was based on the presence of CD45, expressed in inflammatory and hematopoietic cells and F4/80, a specific macrophage surface marker. CD86 and CD206 were used to determine M1 and M2 macrophage subpopulations, respectively (38). Identification of the epithelial cell population was based on the presence of the epithelial cell adhesion molecule (EpCAM) and the absence of CD45. The subsequent display of positive CD24 determined injured proximal tubule epithelia (39). Numbers in quadrants indicate cell proportions in percent of cells that co-express both markers.

### Measurements of oxygen consumption rate

Fatty acid oxidation-associated oxygen consumption rate (OCR) (ligated to oxidative phosphorylation) and extracellular acidification rate (ECAR) (associated with lactate production and glycolysis) were studied using the Seahorse Bioscience metabolic analyzer for the XF Cell Mito Stress Test and Agilent Seahorse XF Real-Time ATP Rate Assay according to the manufacturer’s instructions (40). See **Supplemental Methods** for details.

### Immunofluorescence

A quarter piece of each kidney sample was immersed sequentially in 4% neutral buffered formalin for 24 hours, in 30% sucrose in PBS until tissue sank (6-12 h) and embedded in Tissue-Tek^®^ OCT for cryoprotection at −80°C. OCT blocks were cut in serial frozen sections (10μm thickness). These sections were fixed with 4% PFA for 10 min and permeabilized with 0.25% Triton X-100 in PBS for 5 min at room temperature (RT). Next, they were blocked with 1% BSA in PBS for 30 min at RT and incubated for 1 h for staining with specific primary antibodies and fluorochrome-conjugated secondary antibodies for 1h at RT (**Supplemental Table 6**). Nuclei were stained with DAPI (Sigma, St. Louis, MO) for 5 min at RT. The coverslips were mounted on slides using mowiol (Calbiochem). Tissue fluorescence was visualized by a LSM 510 Meta Confocal microscope with a 40X/1.3 oil Plan-Neofluar M27 objective (Zeiss, Oberkochen, Germany).

### Histology and immunohistochemistry

A quarter piece of each kidney sample was immersed in 4% neutral buffered formalin for 24 hours, embedded in paraffin, cut in serial sections (5μm thickness) and stained with hematoxylin and eosin (H&E), Sirius red and periodic acid-schiff (PAS) stainings as described previously (41) Tissue sections for immunostaining were deparafinized through xylene and hydrated through graded ethanol (100%, 96%, 90%, and 70%) and distilled water. Endogenous peroxidase was blocked. We incubated the tissue sections with 4% BSA for 1 h at RT. Primary antibodies to detect CD3 (M7254, 1:100, Dako, Spain) were incubated for 30 minutes at RT. After washing, slides were incubated with goat anti-mouse HRP secondary antibody (“Visualization Reagent HRP” from SK006, Dako, Spain) for 30 min and visualized using the PD-L1 IHC 22C3 pharmDx (SK006, Dako, Spain). To detect Kim-1, primary antibody (AF1817, 1:500, RyD Systems, Minneapolis, MN) were incubated for 30 minutes at room temperature. To detect F4/80, primary antibody (MCA4976, 1:50, BioRad, Hercules, CA) were incubated at 4°C overnight. After washing, slides were incubated with goat anti-rabbit IgG biotin-SP conjugate secondary antibody (31752, 1:200, Invitrogen, Carlsbad, CA) and rabbit anti-rat IgG biotin-SP (31834, 1:200, Thermo Scientific, Rockford, IL), respectively; and visualized using DAB Substrate Kit (ab64238, abcam, Abcam, Cambridge, UK). Tissue sections were revealed using 3,3’-diaminobenzidine (20 μl/ml, DAB, Dako, Spain) and counterstained with Carazzi’s hematoxylin. Slides were mounted with mowiol (Merck Millipore, Germany). See **Supplemental Methods** for histological and immunohistochemical analysis

### Electron microscopy examination

The number of mitochondria and their structure was analyzed by standard transmission electron microscopy. Renal cortex pieces were cut into small blocks (1 mm^3^) and fixed by immersion in fixative (4% paraformaldehyde/2% glutaraldehyde in 0.1 M phosphate buffer) overnight at 4°C. After washing with cold PBS and incubated in 1% osmium tetroxide and 1% potassium ferricyanide in water for 1 hour at 4°C. Pieces were sequentially stained with 0.15% tannic acid (in 0.1 M phosphate buffer) for 1 min at room temperature (RT), and 2% uranyl acetate in water for 1 hour at RT. Next, they were dehydrated in graded ethanol, and embedded in EmBed812 resin (Electron Microscopy Sciences, Fort Washington, PA). Serial ultrathin sections (70 nm) of the tissue were collected on copper mesh grids. Three grids from each sample were examined using a JEOL 1230 transmission electron microscope, and digital photographs were captured by real-time digital imaging.

### Adenovirus-mediated CPT1A overexpression

CPT1A overexpression in cells was driven and transduced by adenoviral particles. Adenoviruses carrying CPT1A (AdCPT1A) or adenoviruses control (AdControl) were amplified, purified and titrated according to (42) (see **Supplemental Methods** for details). HCK8 cells were seeded into 60 mm culture dishes to reach a confluence of 70%. They were infected with adenoviruses AdControl and AdCPT1A [100 moi (multiplicity of infection)] for 24 h in serum-free DMEM/F12 (supplemented with 15 mM Hepes, 5% FBS, 1x ITS, 0.5 μg/ml hydrocortisone, 50 units/mL penicillin and 50 μg/mL streptomycin) and then the medium was replaced with complete medium for additional 24 h. Adenovirus infection efficiency was assessed in AdControl-infected cells by immunofluorescence.

### Clinical data

Acyl-carnitines were evaluated based on the PREDIMED trial. The PREDIMED is a primary prevention multicenter trial conducted in Spain. A detailed description of this trial is described elsewhere (13). Liquid chromatography–tandem mass spectrometry was used to semi-quantitatively profile acyl-carnitines in plasma samples. Our analysis is based on a subsample of 686 participants with metabolomics data and measures of kidney function including glomerular filtration rate and urinary albumin-creatinine ratio (43, 44). Studies of CPT1A expression in kidney tubules were performed in a cohort of 433 patients (described in (45) and hereon named CKD cohort), whose demographic and clinical features are summarized in **Supplemental table 9**.

### Statistical analysis

Data in experimental models were analyzed using nonparametric tests except where indicated. The difference between two independent groups was examined with Mann-Whitney test, while more than two groups were compared with Kruskal-Wallis test. A P-value of 0.05 or less was considered statistically significant (*^/#^: P < 0.05, * *^/##^: P < 0.01, ^***/###^: P < 0.001). Data were analyzed using GraphPad Prism 6.0 (GraphPad Software, La Jolla, CA). Data are reported as mean ± standard error of mean (SEM). See **Supplemental Material** for statistics of data in clinical studies for acyl-carnitines and CPT1A expression.

### Study approval

Animals were handled in agreement with the Guide for the Care and Use of Laboratory Animals contained in Directive 2010/63/EU of the European Parliament. Approval was granted by the local ethics review board of the Centro de Biología Molecular “Severo Ochoa” in Madrid, the Ethics committee at CSIC and the Regulatory Unit for Animal Experimental Procedures from the Comunidad de Madrid. For human studies informed consent of participants is specified in the references (44, 45).

## Author contributions

SL conceived and directed research. VM designed, performed and analyzed the majority of experiments. JT and LME performed experiments. JIH provided technical assistance for mouse experiments. CC performed histological evaluation. LH and DS provided the CPT1A adenoviruses. MRO, DRP, PC and CB provided intellectual and technical insight. RR assisted with the Taqman analysis. SO coordinated the generation of the CPT1A KI mouse model. XS, KS, MRC, JSS and MAMG performed studies in two different cohorts of CKD patients. All authors helped with the discussion of the results and SL and VM wrote the manuscript.

## Acknowledgements

This work was supported by Grants from the Ministerio de Economía y Competitividad (MINECO) SAF2012-31388 (SL), SAF2015-66107-R (SL), PID2019-104233RB-100 (SL), PI17/01513 (DRP), SAF2017-83813-C3-1-R (DS and LH) co-funded by the European Regional Development Fund, Instituto de Salud Carlos III REDinREN RD12/0021/0009 and RD16/0009/0016 (SL, DRP and MRO), PI17/00119 (MRO), FIS PI17/00625 (DRP), Comunidad de Madrid “NOVELREN” B2017/BMD-3751 (SL, DRP and MRO), a grant-in-aid from the Spanish Society of Nephrology (Fundación Senefro 2017) to SL and Fundación Renal “Iñigo Alvarez de Toledo” (SL), the Centro de Investigación Biomédica en Red de Fisiopatología de la Obesidad y la Nutrición (CIBEROBN) CB06/03/0001 (DS), the Government of Catalonia 2017SGR278 (DS) and the Fundació La Marató de TV3 201627-30 (DS), all from Spain. The CBMSO receives institutional support from Fundación “Ramón Areces”. The Prevención con Dieta Mediterránea (PREDIMED) trial was supported by grants from the agency for biomedical research of the Spanish government, the Instituto de Salud Carlos III RTIC G03/140, the Centro de Investigación Biomédica en Red de Fisiopatología de la Obesidad y Nutrición, the Centro Nacional de Investigaciones Cardiovasculares CNIC 06/2007, the Fondo de Investigación Sanitaria Fondo Europeo de Desarrollo Regional PI04-2239, PI 05/2584, CP06/00100, PI07/0240, PI07/1138, PI07/0954, PI 07/0473, PI10/01407, PI10/02658, PI11/01647, P11/02505, and PI13/00462, the Ministerio de Ciencia e Innovación AGL-2009–13906-C02 and AGL2010–22319-C03, the Fundación Mapfre 2010, Consejería de Salud de la Junta de Andalucía PI0105/2007, the Public Health Division of the Department of Health of the Autonomous Government of Catalonia, Generalitat Valenciana ACOMP06109, GVA-COMP2010–181, GVACOMP2011–151, CS2010-AP-111, and CS2011-AP-042 and the Regional Government of Navarra P27/2011. Metabolomics measurements in PREDIMED were funded by the National Institutes of Health R01DK102896, F31DK114938, NIH/NHLBI 1R01HL118264, NIH/NHLBI 2R01HL118264. PREDIMED funding is related to MRC, MA-MG and JS. VM was supported by pre-doctoral fellowships of the FPI Program (BES-2013-065986 and BES-2014-068929) from MINECO. We acknowledge The J. David Gladstone Institutes and Dr. Ken Nakamura (University of California) for providing the Clover-mApple ATP sensor, Dr. Miguel López (Universidad de Santiago de Compostela) for providing the p-AMPK and p-ACC antibodies, Dr. Maria Luisa Toribio (CBMSO) for her assistance with the flow cytometry analysis and Dr. Germán Andrés (CBMSO) for his help with electron microscopy images evaluation. We are grateful to the laboratories of Jorgina Satrústegui and José Manuel Cuezva for sharing the Seahorse equipment and Laura Santos for her help with histology procedures, from the laboratory of Marta Ruiz-Ortega at the Fundación Jiménez Díaz. We also thank the help of the following facilities of the CBMSO: animal housing, flow cytometry and confocal and electron microscopy.

## Notes

**Conflict of interest:** The authors have declared that no conflict of interest exists.

### Competing Interest Statement

The authors have declared no competing interest.

### Summary of Updates

Figure 7 has been modified.

