## Supplemental material for "Renal tubule Cpt1a overexpression protects from kidney fibrosis by restoring mitochondrial homeostasis"

### Supplemental Figure Legends

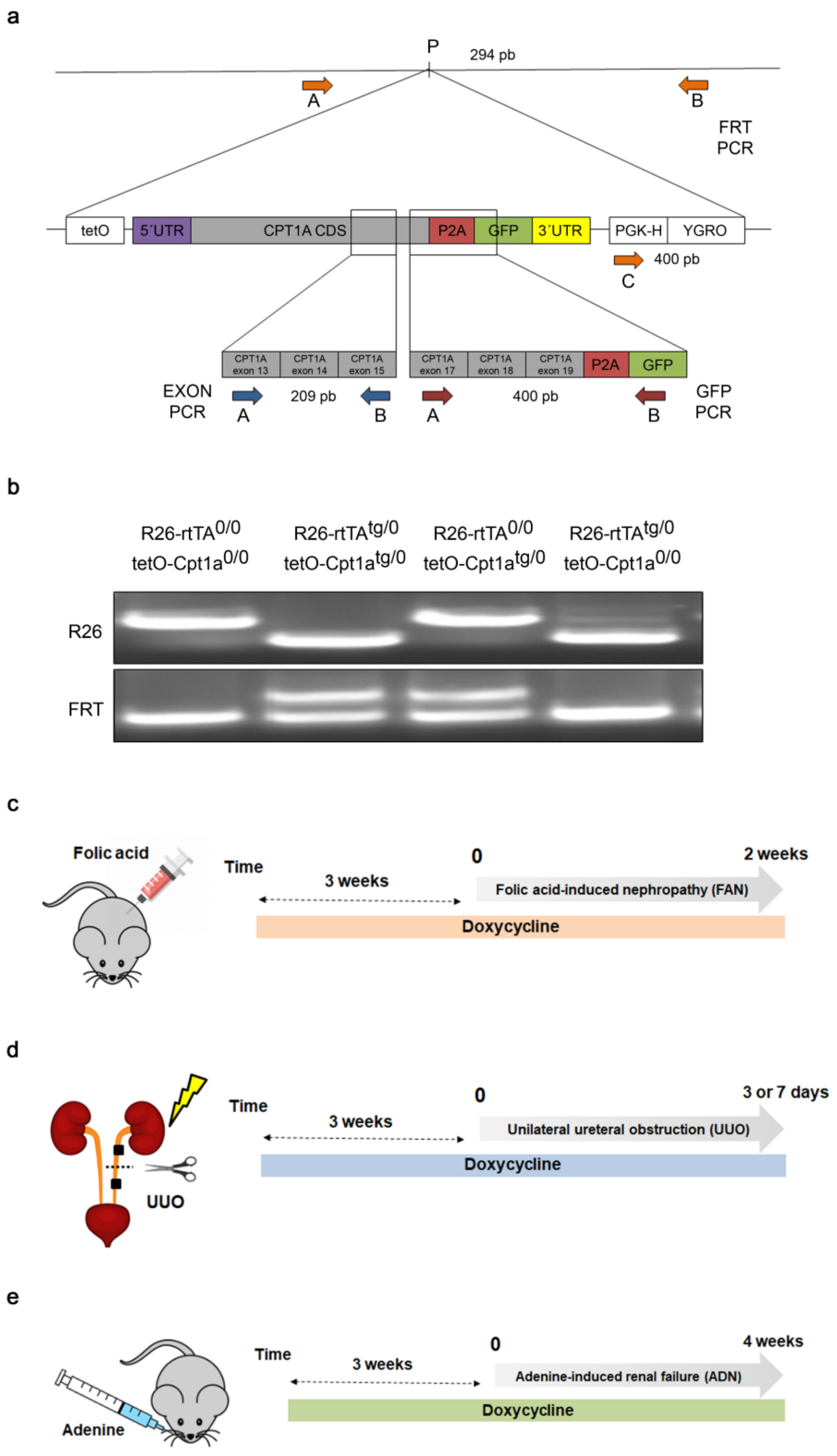


Supplemental Figure 1

**Supplemental Figure 1. Genotypic characterization of the Pax8-rtTA$^{tg/0}$:tetO-Cpt1a$^{tg/0}$ mice and schematics of kidney fibrosis models. (A)** Genetic validation of tetO-Cpt1a$^{tg/0}$ mice. Schematic of the wild-type Col1a1 allele (top) and targeted with the *Cpt1a* gene construction (expanded). Genomic DNA analysis for the CPT1A$^{tetO}$ allele was performed with 3 different PCR strategies, named as referred in Table 1: FRT, Exon and GFP. The positions of primers for these PCR strategies are indicated by colored arrows: FRT, orange; Exon, blue; GFP, red. For FRT PCR, amplicons are 294 bp (AB primers) and 400 bp (CB primers) corresponding to the WT and CPT1A$^{tetO}$ alleles, respectively. For GFP PCR the amplicon is 400 bp. For Exon PCR the amplicon is 209 bp. In both cases they correspond to the CPT1A$^{tetO}$ allele. P: PstI restriction site. **(B)** Genomic DNA analysis by PCR for the Rosa26 allele variants generates 650-bp and 300-bp amplicons for WT (R26-WT) and ROSA26-M2-rtTA (R26-rtTA) alleles, respectively as indicated in **Supplemental Table 1**. The FRT PCR is detailed in **(A)**, the presence of two bands (**lanes 2 and 3**) denotes heterozygosity. Animals harboring CPT1A$^{tetO}$ allele and lacking ROSA26-M2-rtTA allele (**lane 3**) were selected for crossing with Pax8-rtTA$^{tg/0}$ mice. (**C, D, E**) Timeline of the experimental procedures corresponding to folic acid nephropathy-FAN (**C**), unilateral ureteral obstruction-UUO (**D**) and adenine induced nephrotoxicity-ADN (**E**).

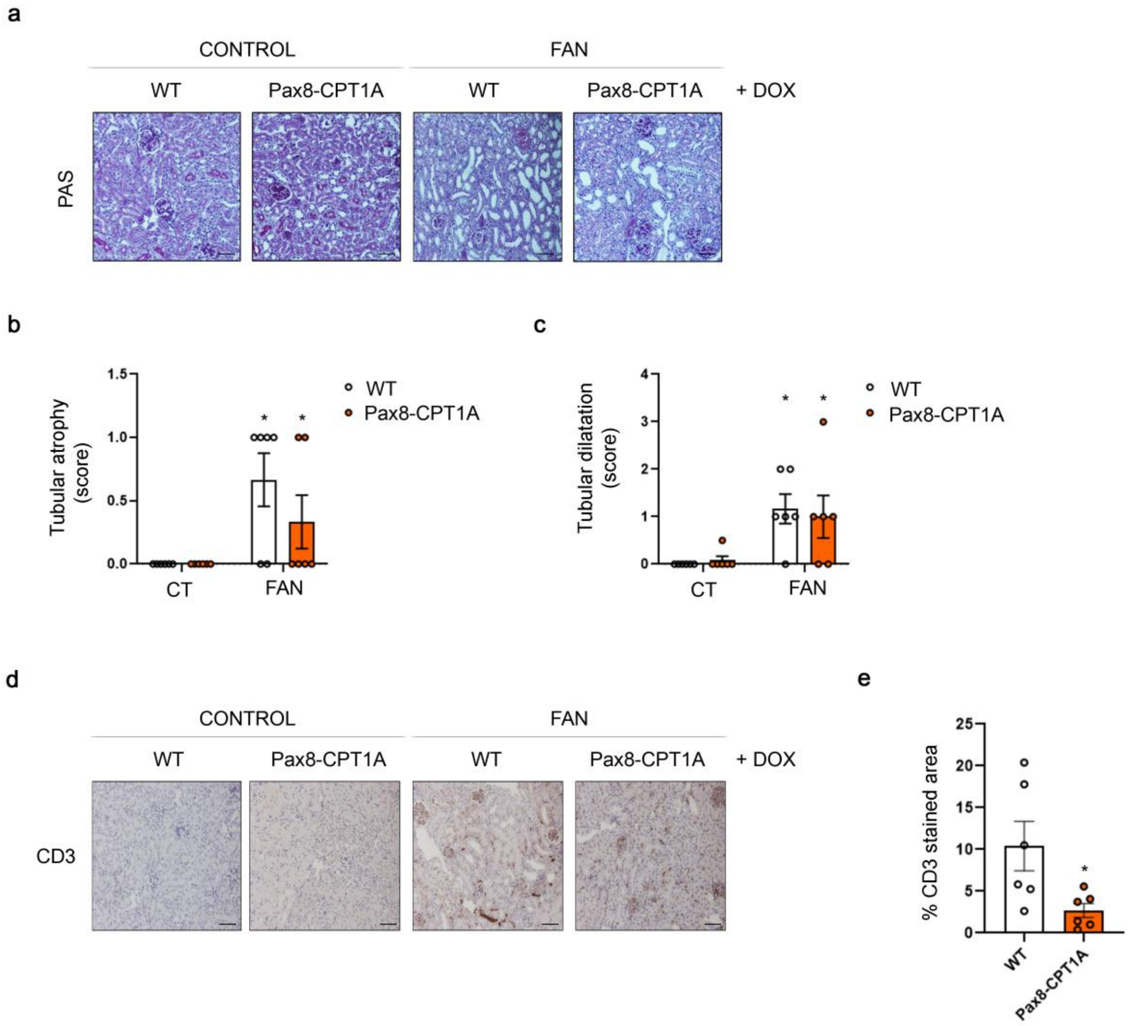


Supplementary Figure 2

**Supplemental Figure 2. Renal tubule induction of CPT1A prevents kidney damage-induced structural tubular alteration and renal T lymphocyte infiltration in the FAN model. (A)** Representative microphotographs from one mouse per group of Periodic Acid Schiff (PAS) staining of kidneys from WT and Pax8-CPT1A mice subjected to FAN after doxycycline treatment (Dox). Scale bars: 50 µm. Semi-quantitative determinations (grade 0 to 4) of tubular atrophy **(B)** and tubular dilatation **(C)** in FAN related to control kidneys. Bar graphs represent the mean ± s.e.m, n = 6 mice. $^*P < 0.05$, compared to control kidneys in WT mice, respectively. **(D)** Representative micrographs of one mouse per group showing the expression of CD3 in kidney sections of mice treated as described above. Scale bar = 50 µm. **(E)** Bar graph represents the quantification of the % of CD3 positive stained area relative in FAN vs WT, mean ± s.e.m, n = 6 mice, $^*P < 0.05$. Statistical significance between two independent groups was determined using non-parametric two-tailed Mann-Whitney test, while more than two groups were compared with Kruskal-Wallis test.

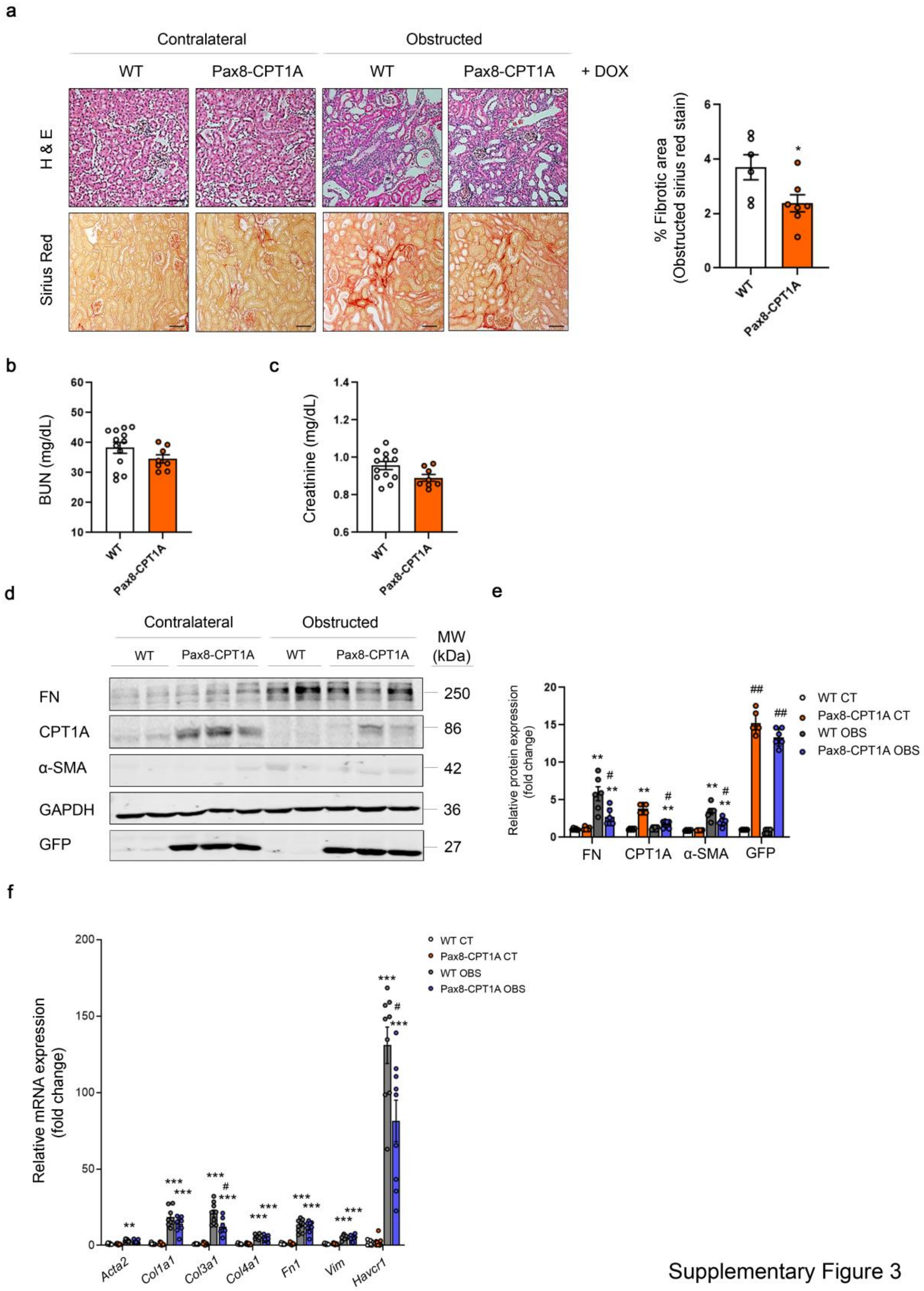


Supplementary Figure 3

**Supplemental Figure 3. Renal tubule overexpression of CPT1A prevents UUO-dependent increased expression of fibrosis-associated markers. (A)** Representative microphotographs from one mouse per group of hematoxylin and eosin (H&E) (upper panels) and Sirius Red (lower panels) staining of kidneys from WT and Pax8-CPT1A mice subjected to UUO for 7 days after doxycycline treatment (Dox). Scale bars: 50 µm. Quantification of Sirius Red staining represents the mean ± s.e.m, n = 6 mice per group. *P < 0.05, compared to obstructed kidneys in WT mice, respectively. **(B)** Serum blood urea nitrogen (BUN) levels of WT and Pax8-CPT1A mice subjected to UUO for 7 days after doxycycline treatment. Data represent the mean ± s.e.m (at least n = 8 mice). **(C)** Serum creatinine levels of WT and Pax8-CPT1A mice subjected to UUO for 7 days after doxycycline treatment. Data represent the mean ± s.e.m (at least n = 8 mice). **(D)** Immunoblots depicting fibronectin (FN), carnitine palmitoyltransferase 1A (CPT1A), alpha-smooth muscle actin (α-SMA), GAPDH and green fluorescence protein (GFP) protein levels in contralateral (CT) and obstructed (OBS) kidneys from 3 WT and 3 Pax8-CPT1A mice. **(E)** Bar graphs represent the mean ± s.e.m. of fold changes corresponding to densitometric analyses (n = 6 mice). **P < 0.01 compared to their corresponding contralateral (CT) kidneys; $^{\#}$P < 0.05, $^{\#\#}$P < 0.01 compared to kidneys from WT mice with the same experimental condition. **(F)** mRNA levels of fibrosis-associated genes were determined by qRT-PCR using TaqMan qPCR probes in contralateral (CT) and obstructed (OBS) kidneys from WT and Pax8-CPT1A mice subjected to UUO for 7 days after doxycycline induction. Bar graphs represent the mean ± s.e.m. of fold changes (n = 9 mice). ***P < 0.01 compared to their corresponding contralateral (CT) kidneys; $^{\#}$P < 0.05 compared to kidneys from WT mice with the same experimental condition. Statistical significance between two independent groups was determined using non-parametric two-tailed Mann-Whitney test, while more than two groups were compared with Kruskal-Wallis test. For detailed gene nomenclature see **Supplemental Table 4**.

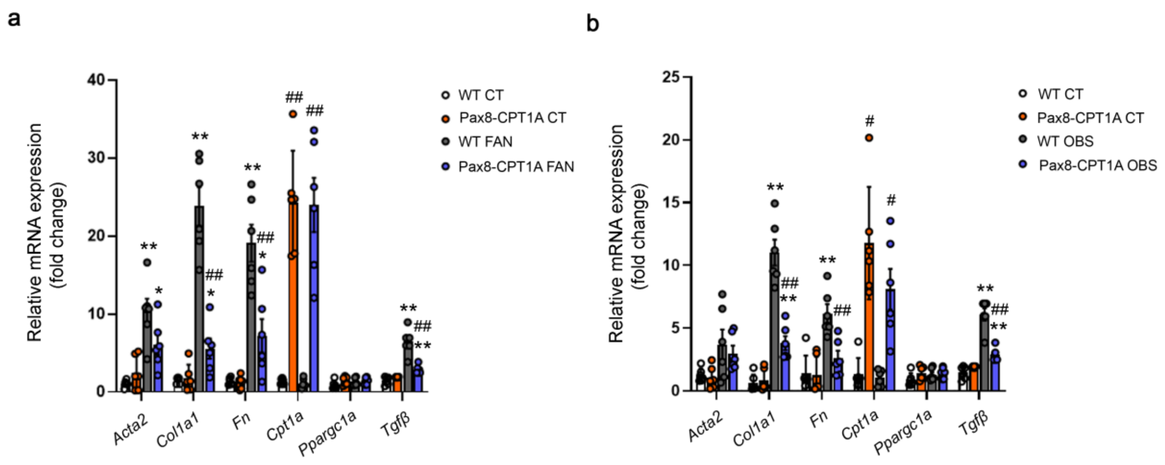


Supplementary Figure 4

**Supplemental Figure 4. Renal tubule induction of CPT1A prevents kidney damage-induced expression of fibrosis-related genes. (A, B)** mRNA levels of alpha-smooth muscle actin (α-SMA), alpha 1 type-1 collagen (Col1α1), fibronectin (FN), carnitine palmitoyltransferase 1A (CPT1A), peroxisome proliferator-activated receptor gamma coactivator 1 alpha (Ppargc1a) and transforming growth factor beta (TGF-β) were determined by qRT-PCR using Sybr green in kidneys of WT and Pax8-CPT1A mice subjected to FAN (A) or UUO (B) after doxycycline induction. Bar graphs represent the mean ± s.e.m. of fold changes (n = 6 mice). *P < 0.05, **P < 0.01 compared to their corresponding control (CT) kidneys; $^{\#}$P < 0.05, $^{\#\#}$P < 0.01, compared to kidneys from WT mice with the same experimental condition. Statistical significance between two independent groups was determined using non-parametric two-tailed Kruskal-Wallis test.

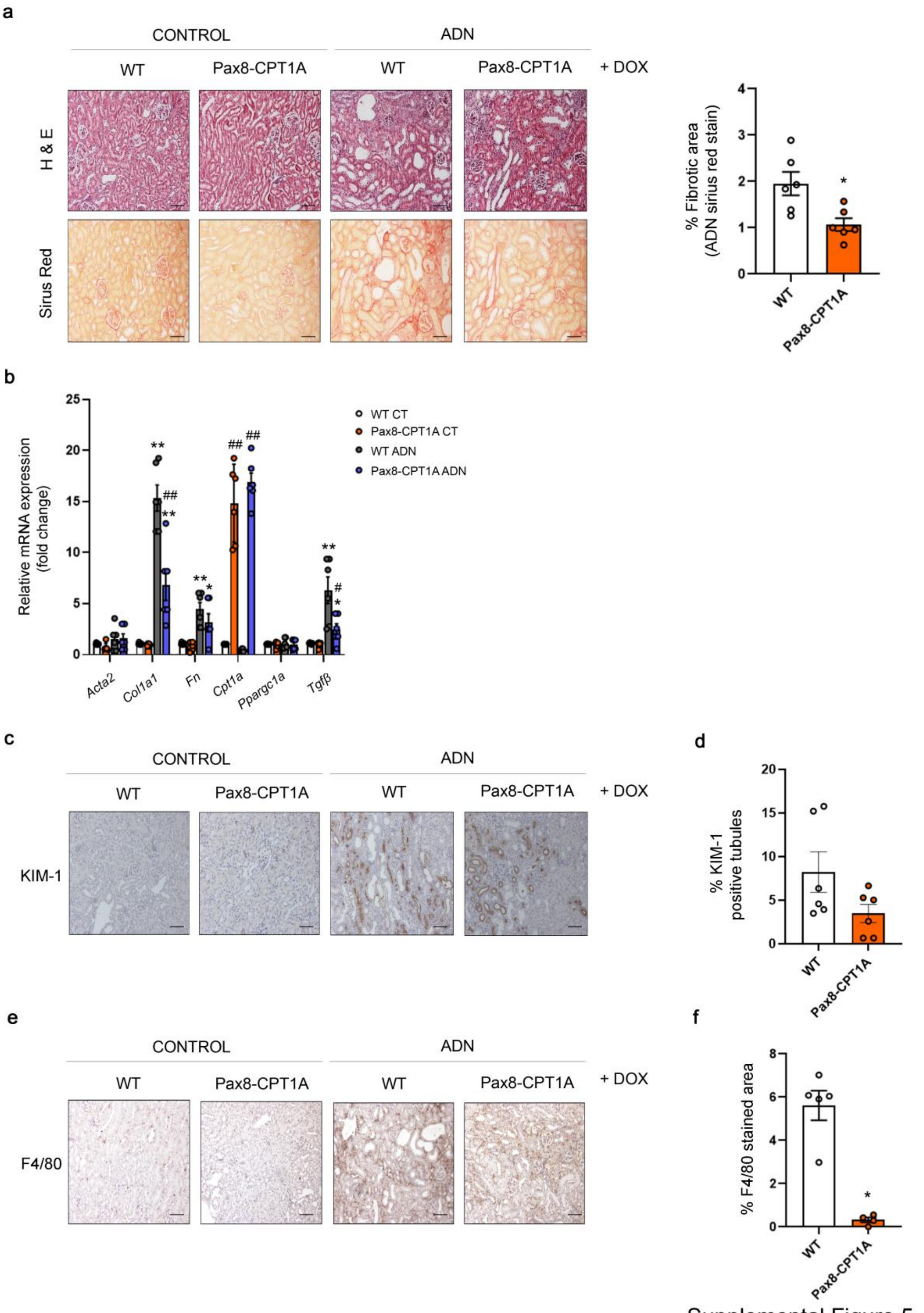


Supplemental Figure 5

**Supplemental Figure 5. Renal tubule overexpression of CPT1A prevents ADN-dependent experimental renal fibrosis. (A)** Representative microphotographs from one mouse per group of hematoxylin and eosin (H&E) (upper panels) and Sirius Red (lower panels) staining of kidneys from WT and Pax8-CPT1A mice subjected to ADN after doxycycline treatment (Dox). Scale bars: 50 µm. Quantification of Sirius Red staining represents the mean ± s.e.m, n = 6 mice per group. *P < 0.05, compared to ADN kidneys in WT mice, respectively. **(B)** mRNA levels of alpha-smooth muscle actin (α-SMA), alpha 1 type-1 collagen (Col1α1), fibronectin (FN), carnitine palmitoyltransferase 1A (CPT1A), peroxisome proliferator-activated receptor gamma coactivator 1 alpha (Ppargc1a) and transforming growth factor beta (TGF-β) were determined by qRT-PCR using Sybr green in kidneys of WT and Pax8-CPT1A mice subjected to adenine-induced nephrotoxicity (ADN) after doxycycline induction. Bar graphs represent the mean ± s.e.m. of fold changes (n = 6 mice). *P < 0.05, **P < 0.01 compared to their corresponding control (CT) kidneys; $^{\#}$P < 0.05, $^{\#\#}$P < 0.01, compared to kidneys from WT mice with the same experimental condition. **(C, E)** Representative micrographs of one mouse per group showing the expression of KIM-1 **(C)** and F4/80 **(E)** in kidney sections of mice treated as described above. Scale bar = 50 µm. **(D, F)** Bar graph represents the quantification of the % of KIM-1 **(D)** and F4/80 **(F)** positive stained area in adenine-treated mouse kidneys (ADN). Bar graphs represent the mean ± s.e.m, n = 6 mice. *P < 0.05, compared to ADN kidneys in WT mice, respectively. Statistical significance between two independent groups was determined using non-parametric two-tailed Mann-Whitney test, while more than two groups were compared with Kruskal-Wallis test.

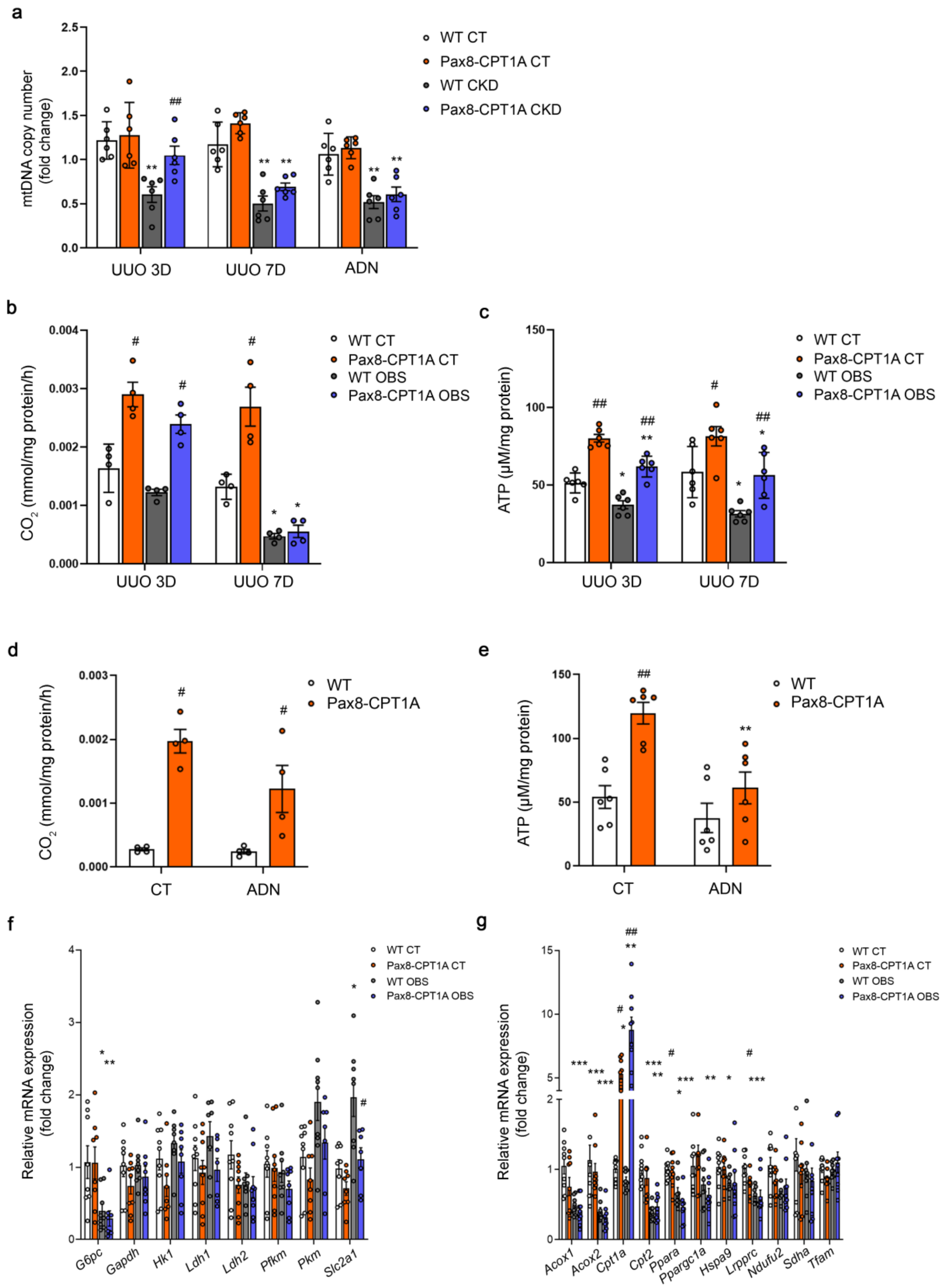


Supplemental Figure 6

**Supplemental Figure 6. CPT1A overexpression mitigates mitochondrial and impaired fatty acid oxidation induced by UUO. (A)** Mitochondrial DNA copy number (mtDNA) was determined in kidneys of WT and Pax8-CPT1A mice in the 3 and 7-days UUO and adenine-induced nephrotoxicity (ADN) models after doxycycline induction. (**B, D**) Radiolabeled palmitate-derived $CO_2$ was determined after incubation of $^{14}C$-palmitate with kidney tissue from WT and Pax8-CPT1A mice in the 3 and 7-days UUO **(B)** and adenine-induced nephrotoxicity (ADN) **(D)** models after doxycycline induction. **(C, E)** ATP levels in total kidney tissue determined in mice subjected to 3 and 7-days UUO **(C)** or ADN **(E)**. Bar graphs represent the mean ± s.e.m (n = 4 mice in **B** and **D**, n = 6 mice in **A**, **C** and **E**). *P < 0.05, **P < 0.01 compared to their corresponding control (CT) kidneys; $^{\#}$P < 0.05, $^{\#\#}$P < 0.01 compared to kidneys from WT mice with the same experimental condition. **(F, G)** mRNA levels of glucose utilization- **(F)** and peroxisomal/mitochondrial function- **(G)** associated genes were determined by qRT-PCR using TaqMan qPCR probes in contralateral (CT) and obstructed (OBS) from kidneys of WT and Pax8-CPT1A mice subjected to UUO for 7 days after doxycycline induction. Bar graphs represent the mean ± s.e.m. of fold changes (n = 9 mice). *P < 0.05, **P < 0.01, ***P < 0.001 compared to their corresponding contralateral (CT) kidneys; $^{\#}$P < 0.05, $^{\#\#}$P < 0.01 compared to kidneys from WT mice with the same experimental condition. Statistical significance between two independent groups was determined using non-parametric two-tailed Kruskal-Wallis test. For detailed gene nomenclature see **Supplemental Table 4**.

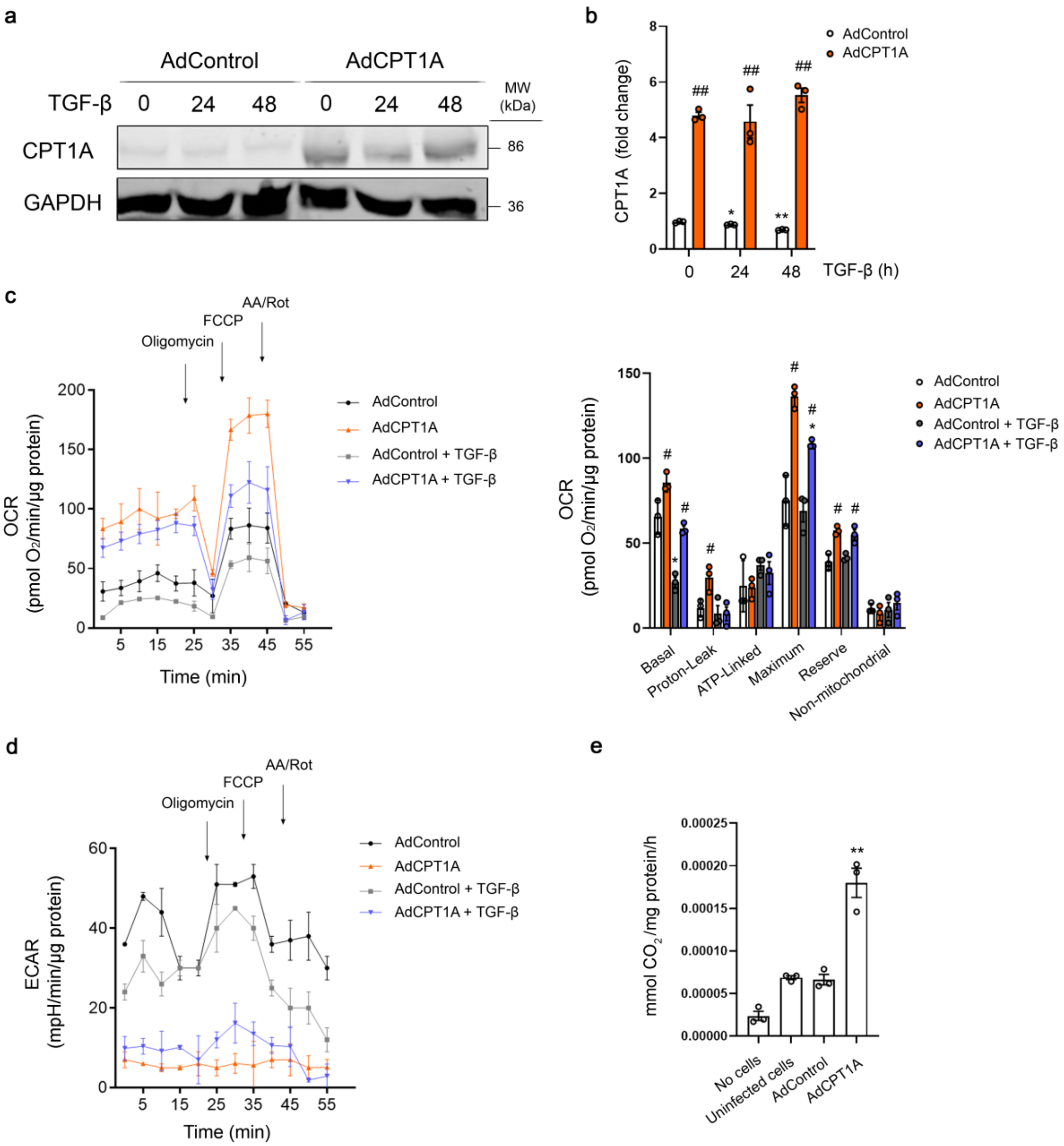


f

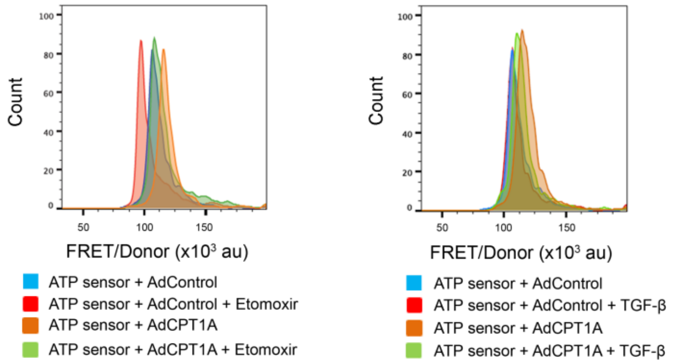


| | Median (FRET/Donor) ($x10^3$) |
|---|---|
| ATP sensor + AdControl | 97±1 |
| ATP sensor + AdControl + Etomoxir | 84±1.6 # |
| ATP sensor + AdCPT1A | 128±0.8 * |
| ATP sensor + AdCPT1A + Etomoxir | 110±0.4 # |

| | Median (FRET/Donor) ($x10^3$) |
|---|---|
| ATP sensor + AdControl | 106±0.2 |
| ATP sensor + AdControl + TGF-β | 95±1.1 # |
| ATP sensor + AdCPT1A | 125±1.5 * |
| ATP sensor + AdCPT1A + TGF-β | 121±0.01 |

Supplemental Figure 7

**Supplemental Figure 7. HKC-8 cells transduced with *CPT1A* exhibit reduced TGF-β1-induced FAO inhibition and fibrogenic transformation. (A)** Immunoblot depicts CPT1A protein levels in HKC-8 cells infected with adenoviruses carrying CPT1A (AdCPT1A) or AdControl and exposed to 10 ng/ml TGF-β1 for the indicated time points. **(B)** Bar graphs represent the mean ± s.e.m. of fold changes corresponding to densitometric analyses (n = 3 independent experiments). *P < 0.05, **P < 0.01 compared to their corresponding untreated cells; $^{\#\#}$P < 0.01 compared to cells bearing AdControl with the same experimental treatment. **(C)** Oxygen consumption rate (OCR) of HCK-8 cells expressing AdCPT1A or its negative control AdControl was measured with a Seahorse XF24 Extracellular Flux Analyser. Bar graphs (right panel) show the rates of OCR associated to basal, proton-leak, ATP-linked, maximum reserve capacity and non-mitochondrial respiratory statuses. Each data point represents the mean ± s.e.m of 4 independent experiments, each performed in triplicate. *P < 0.05 compared to their corresponding control (CT) cells; $^{\#}$P < 0.05 compared to cells bearing AdControl with the same experimental treatment. **(D)** Extracellular acidification rate (ECAR) of cells treated as in (C). Data are normalized by protein amount. Each data point represents the mean ± s.e.m of 4 independent experiments, each performed in triplicate. (**E**) Radiolabeled palmitate-derived $CO_2$ was determined after incubation of cells treated as in (A) with $^{14}$C-palmitate. **P < 0.01 compared to their corresponding cells bearing AdControl. Each data point represents the mean ± s.e.m of 4 independent experiments, each performed in triplicate. **P < 0.05 compared to cells bearing AdControl with the same experimental treatment. **(F)** FRET/donor distributions of HKC-8 cells containing the Clover-mApple ATP sensor and expressing AdCPT1A or AdControl. Where indicated, cells were treated with etomoxir or TGF-β. Tables (right panel) show fluorescence intensity of FRET/Donor ratio, median ± s.e.m of 4 independent experiments, each performed in triplicate. *P < 0.05 compared to their corresponding control (CT) cells; $^{\#}$P < 0.05 compared to cells bearing AdControl with the same experimental treatment. Statistical significance between two independent groups was determined using non-parametric two-tailed Kruskal-Wallis test.

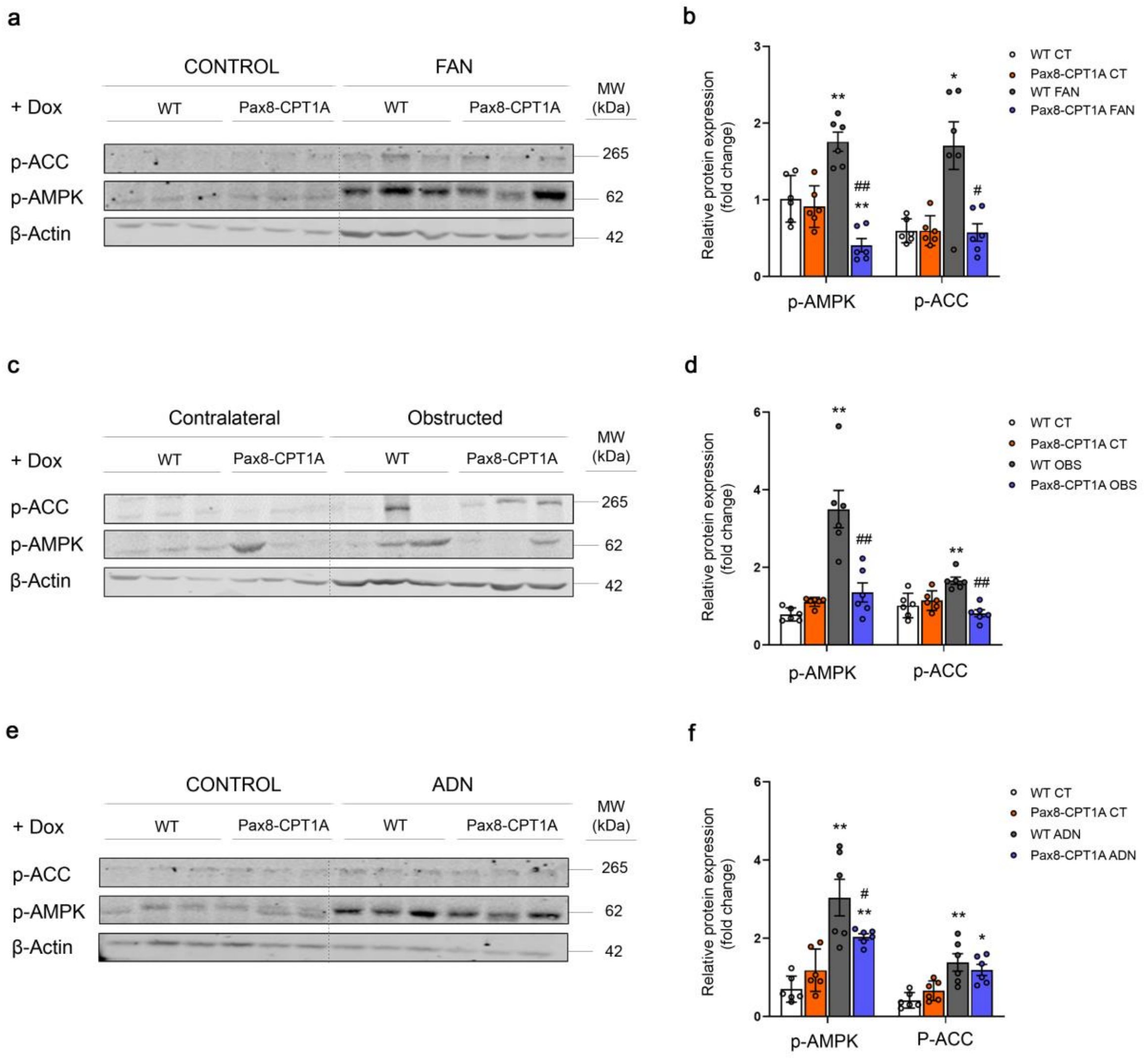


Supplemental Figure 8

**Supplemental Figure 8. Genetic CPT1A overexpression prevents AMP-activated protein kinase (AMPK) activation associated to kidney fibrosis.** Representative immunoblots corresponding to phosphorylated AMP-activated protein kinase (p-AMPK) and acetyl-CoA carboxylase (p-ACC) protein levels in kidneys of WT and Pax8-CPT1A mice subjected to FAN **(A)**, UUO **(C)** or adenine-induced nephrotoxicity (ADN) **(E)** after doxycycline induction (represented are 3 mice). **(B, D, F)** Bar graphs represent the mean ± s.e.m. of fold changes corresponding to densitometric analyses of immunoblots corresponding to the FAN **(B)**, UUO **(D)** and ADN **(F)** model (n = 6 mice per group). β-actin was used for normalization purposes. *P < 0.05, **P < 0.01 compared to their corresponding control (CT) kidneys; $^{\#}$P < 0.05, $^{\#\#}$P < 0.01 compared to kidneys from WT mice with the same experimental condition. Statistical significance between two independent groups was determined using non-parametric two-tailed Kruskal-Wallis test.

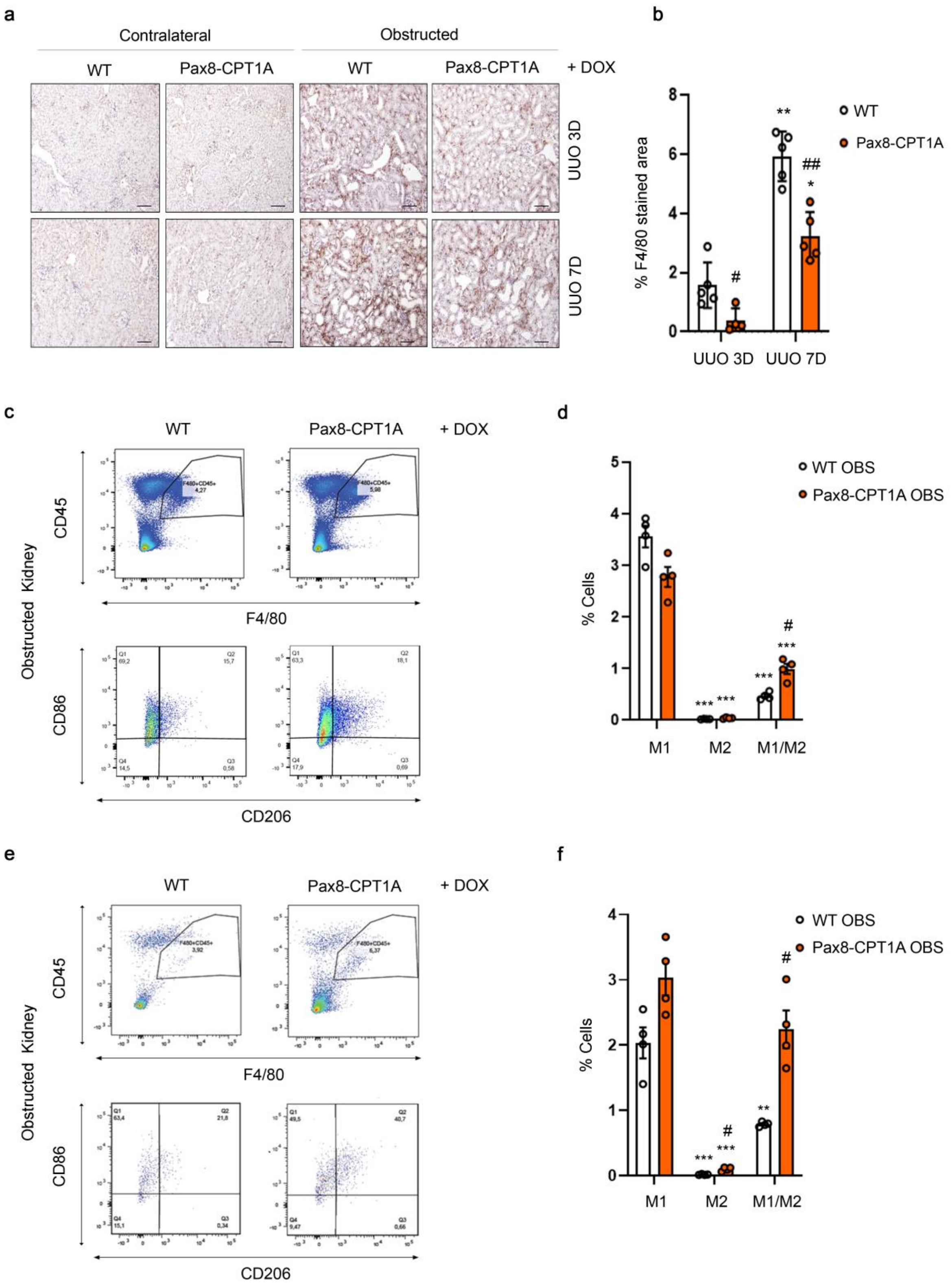


Supplementary Figure 9

**Supplemental Figure 9. Overexpression of CPT1A increases the infiltration of macrophage subpopulations expressing CD206 (M2) and both CD86 and CD206 (M1/M2) in the 7-days UUO model. (A)** Representative micrographs of one mouse per group showing the expression of F4/80 in kidney sections of mice treated as described above. Scale bar = 50 µm. **(B)** Bar graph represents the quantification of the % of F4/80 positive stained area in obstructed kidneys from WT and CPT1A KI mice subjected to UUO for 3 (UUO 3D) and 7 days (UUO 7D) after doxycycline induction. Bar graphs represent the mean ± s.e.m, n = 6 mice. *P < 0.05, **P < 0.01, compared to obstructed kidneys in WT mice subjected to UUO for 3 days (UUO 3D). $^{\#}$P < 0.05, $^{\#\#}$P < 0.01 compared to kidneys from WT mice with the same experimental condition. **(C, E)** Representative multiparameter flow cytometry dot plots showing the expression of macrophage populations, studied as in **Figure 5**, in kidney cells from WT and Pax8-CPT1A mice subjected to 3 **(C)** and 7 **(E)** -days UUO after doxycycline induction (upper panels) (one mouse per group is represented). CD86 and CD206 were used to determine the proportion of M1 and M2 macrophage subpopulations, respectively, in the total macrophage population (F4/80+, CD45+) (lower panels). Numbers in quadrants indicate cell proportions in percent of cells that express both markers in **C** and **E**, respectively. **(D, F)** Bar graph represents the percentage of kidney cells expressing CD86 (M1), CD206 (M2) or both (M1/M2) markers. Data represent mean ± s.e.m (n = 4 mice per group). **P < 0.01, ***P < 0.001 compared to % of M1 subpopulation in obstructed kidneys from WT mice; $^{\#}$P < 0.05 compared to corresponding cell subpopulation in obstructed kidneys from WT mice. Statistical significance between two independent groups was determined using non-parametric two-tailed Kruskal-Wallis test.

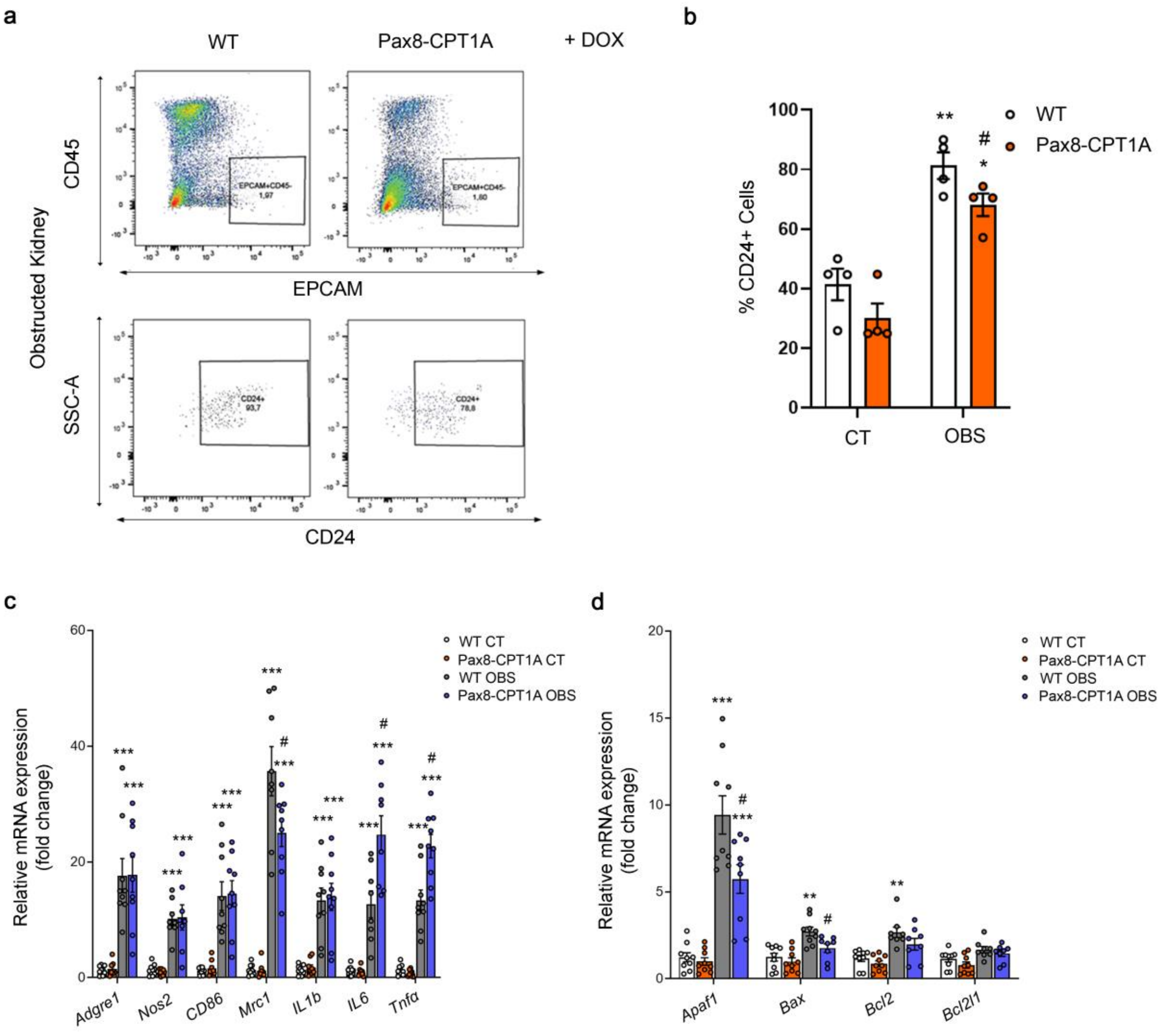


Supplementary Figure 10

**Supplemental Figure 10. CPT1A overexpression reduces epithelial cell damage in the UUO model. (A)** Representative flow cytometry dot plots illustrating cell populations from obstructed kidneys of WT and Pax8-CPT1A mice subjected to UUO for 7 days after doxycycline treatment. Cells were gated for CD45 negative (-)/Epithelial Cell Adhesion Molecule (EpCAM) positive (+) (upper panels) and then selected for the presence of CD24 (lower panels). Numbers in quadrants indicate cell proportions in percent. Data shown are representative of 4 mice per group. **(B)** Bar graphs show the percentage of kidney cells positive for CD24. Data represent the mean ± s.e.m (n = 4 mice). *P < 0.05, **P < 0.01 compared to their corresponding control (CT) kidneys; $^{\#}$P < 0.05 compared to damaged kidneys in WT mice. **(C, D)** mRNA levels of inflammation **(C)** and apoptosis-associated **(D)** genes were determined by qRT-PCR using TaqMan qPCR probes in contralateral (CT) and obstructed (UUO) from kidneys of WT and Pax8-CPT1A mice subjected to UUO for 7 days after doxycycline induction. Bar graphs represent the mean ± s.e.m. of fold changes (n = 9 mice). **P < 0.01, ***P < 0.001 compared to their corresponding contralateral (CT) kidneys; $^{\#}$P < 0.05 compared to kidneys from WT mice with the same experimental condition. Statistical significance between two independent groups was determined using non-parametric two-tailed Kruskal-Wallis test. For detailed gene nomenclature see **Supplemental Table 4**.

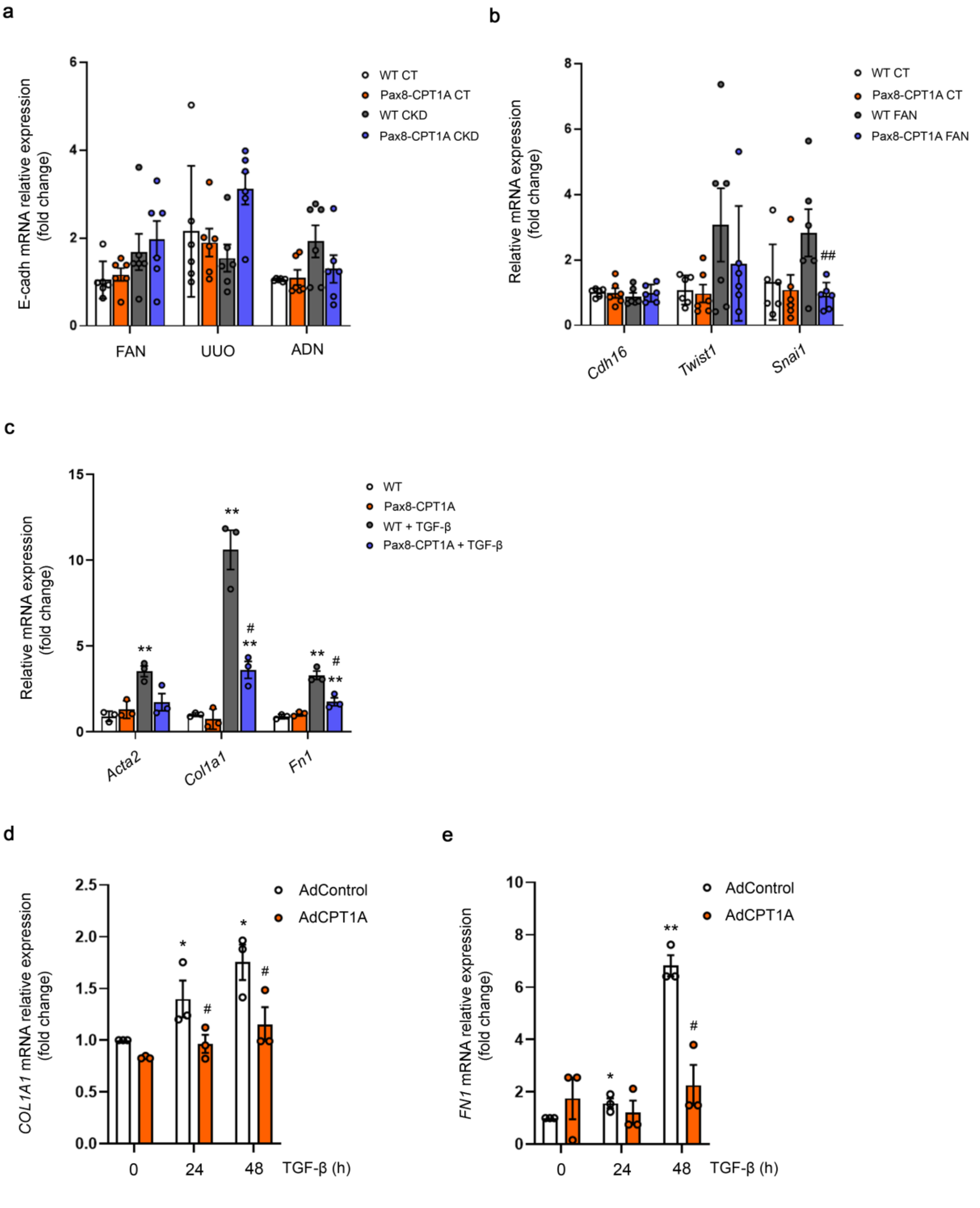


Supplemental Figure 11

**Supplemental Figure 11. CPT1A overexpression reduces epithelial cell dedifferentiation. (A)** mRNA levels of E-cadherin (E-cadh) were determined by qRT-PCR using Sybr green in kidneys of WT and Pax8-CPT1A mice subjected to UUO, FAN or adenine-induced nephrotoxicity (ADN), respectively, after doxycycline induction. Bar graphs represent the mean ± s.e.m. of fold changes (n = 6 mice). **(B)** mRNA levels of EMT-associated genes were determined by qRT-PCR using TaqMan qPCR probes in kidneys from control (CT) and FAN-treated (FAN) WT and Pax8-CPT1A mice after doxycycline induction. Bar graphs represent the mean ± s.e.m. of fold changes (n = 6 mice). $^{##}P < 0.01$ compared to kidneys from WT mice with the same experimental condition. For gene nomenclature see **Supplemental Table 4**. **(C)** mRNA levels of alpha-smooth muscle actin (Acta2), alpha 1 type-1 collagen (Col1α1), fibronectin (Fn1) from cells of kidneys from WT and CPT1A KI mice were determined by qRT-PCR using Sybr green. Cells were treated with TGF-β (10 ng/ml) where indicated. Bar graphs represent the mean ± s.e.m. of fold changes from 3 independent experiments, each performed in triplicate. $^{**}P < 0.01$ compared to their corresponding untreated cells; $^{\#}P < 0.05$ compared to cells from WT mice with the same experimental condition. **(D, E)** mRNA levels of alpha 1 type-1 collagen (COL1A1) **(D)** and fibronectin (FN1) **(E)** in HKC-8 cells infected with adenoviruses carrying CPT1A (AdCPT1A) or AdControl and exposed to TGF-β (10 ng/ml) were determined by qRT-PCR using Sybr green. Bar graphs represent the mean ± s.e.m. of fold changes from 3 independent experiments, each performed in triplicate. $^{*}P < 0.05$, $^{**}P < 0.01$ compared to their corresponding untreated cells; $^{\#}P < 0.05$ compared to cells bearing AdControl with the same experimental treatment. Statistical significance between two independent groups was determined using non-parametric two-tailed Kruskal-Wallis test.

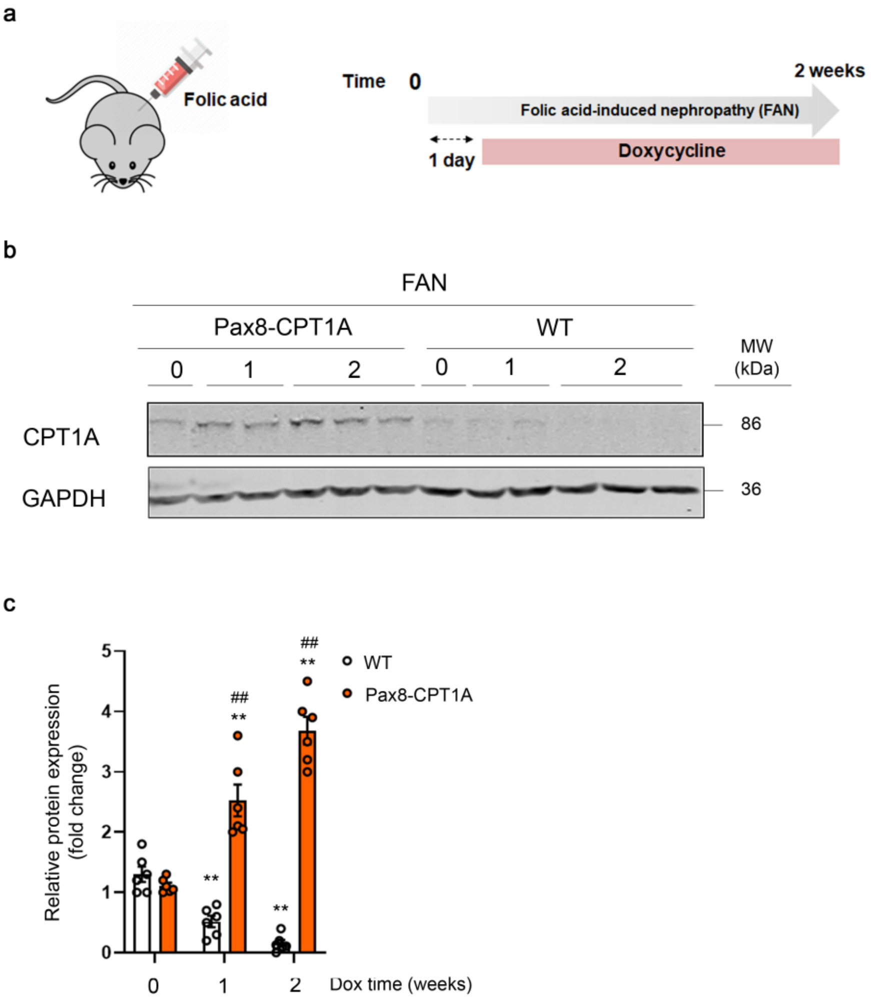


Supplemental Figure 12

**Supplemental figure 12. Expression of CPT1A after doxycycline in the post-damage FAN model. (A)** Timeline of doxycycline administration and FA injection. **(B)** Immunoblots corresponding to carnitine palmitoyltransferase 1A (CPT1A) in kidneys from FA-treated (FAN) WT and Pax8-CPT1A mice at the indicated time-points after doxycycline administration. **(C)** Bar graphs represent the mean ± s.e.m. of fold changes corresponding to densitometric analyses (n = 6 mice). **P < 0.01 compared to their corresponding kidneys from mice with no doxycycline treatment; ##P < 0.01 compared to kidneys from WT mice with the same experimental condition. Statistical significance between two independent groups was determined using non-parametric two-tailed Mann-Whitney test, while more than two groups were compared with Kruskal-Wallis test.

| PCR name | PCR product | 5´- 3´ primer sequence | PCR Amplicons | PCR protocol |
|---|---|---|---|---|
| **Rosa26-rtTA** | *Rosa26* | RV1: GCGAAGAGTTTGTCCTCAACC<br>RV2: GGAGCGGGAGAAATGGATATG<br>FW: AAAGTCGCTCTGAGTTGTTAT | Mutant allele: 300 bp<br>Wild type allele: 650 bp | Denature: 94º 30'<br>Annealing: 59º 45''<br>Extension: 65º 2'30''<br>40x |
| **Pax8-rtTA** | *Pax8-rtTA* construct | FW: CTGGAGAACGCACTGTACGC<br>RV: CCA ATACGCAGCCCAGTGT | Mutant allele: 450 bp<br>Wild type allele: no band | Denature: 95º 3'<br>Annealing: 95º 5''<br>Extension: 60º 30''<br>40x |
| **FRT** | *Col1a1* | FW1 (A):GCAGAAGCGCGGCCGTCTGG<br>RV (B):CCCTCCATGTGTGACCAAGG<br>FW2 (C): GCACAGCATTGCGGACATGC | Mutant allele: 450 bp<br>Wild type allele: 300 pb | Denature: 94º 30'<br>Annealing: 59º 45''<br>Extension: 65º 2'30''<br>40x |
| **GFP** | *Col1a1* | FW (A): AGCAGCAGGTGGAACTGTTT<br>RV (B):CTGAACTTGTGGCCGTTTAC | Mutant allele: 400 bp<br>Wild type allele: no band | Denature: 94º 30'<br>Annealing: 59º 45''<br>Extension: 65º 2'30''<br>40x |
| **Exon** | *Col1a1* | FW (A): CCAGGCTACAGTGGGACATT<br>RV (B):GAACTTGCCCATGTCCTTGT | Mutant allele: 209 bp<br>Wild type allele: no band | Denature: 94º 30'<br>Annealing: 59º 45''<br>Extension: 65º 2'30''<br>40x |

### Supplemental Tables and Supplemental Table Legends

**Supplemental Table 1.** Primer sequences for CPT1A KI mice genotyping.

| Protein | Species | Reference | Dilution | Experimental conditions |
|---|---|---|---|---|
| **CPT1A** | Mouse | Ab128568, Abcam | 1:1000 | 4ºC O/N |
| **α-tubulin** | Mouse | T9026, Sigma-Aldrich | 1:5000 | 1h RT |
| **β-actin** | Mouse | A1978, Sigma-Aldrich | 1:15000 | 1h RT |
| **GFP** | Mouse | AB10145, Sigma-Aldrich | 1:5000 | 4ºC O/N |
| **FN1** | Rabbit | F7387, Sigma, | 1:1000 | 4ºC O/N |
| **α-SMA** | Mouse | sc-32251,<br>Santa Cruz Biotechnology | 1:1000 | 4ºC O/N |
| **GAPDH** | Mouse | MAB374, Millipore | 1:15000 | 1h RT |
| **p-ACC-Ser79** | Rabbit | 07-303, Sigma-Aldrich | 1:1000 | 4ºC O/N |
| **p-AMPKα-Thr172** | Rabbit | 2535, Cell signaling | 1:1000 | 4ºC O/N |
| **COL III** | Mouse | Ab6310, Abcam | 1:1000 | 4ºC O/N |
| **p-RIPK3** | Rabbit | Ab19517, Abcam | 1:1000 | 4ºC O/N |

**Supplemental Table 2.** Primary antibodies for western blot analysis. O/N: overnight, RT: room temperature.

| Gene | 5´- 3´ primer sequence |
|---|---|
| **Mouse** | |
| *Cpt1a* | FW: GGTCTTCTCGGGTCGAAAGC |
| | RV: TCCTCCCACCAGTCACTCAC |
| *Acta2* | FW: CTGACAGAGGCACCACTGAA |
| | RV: CATCTCCAGAGTCCAGCACA |
| *Col1a1* | FW: CTGCTGGCAAAGATGGAGA |
| | RV: ACCAGGAAGACCCTGGAATC |
| *Fn1* | FW: ACCGACAGTGGTGTGGTCTA |
| | RV: CACCATAAGTCTGGGTCACG |
| *Cdh1* | FW: ACCTCCGTGATGAAGGTCTC |
| | RV: CCGGTGTCCCTATTGACAGT |
| *Ppargc1a* | FW: CCCTGCCATTGTTAAGACC |
| | RV: TGCTGCTGTTCCTGTTTTC |
| *Tgfb1* | FW: AGCGGACTACTATGCTAAAGAGGTCAACCC |
| | RV: CCAAGGTAACGCCAGGAATTGTTGCTATA |
| *Ripk1* | RV: TTCGGGAAGGTGTCCTTGTG |
| | FW: CATTGTACTCAGCGCGGTTG |
| *Ripk3* | RV: CTCCGTGCCTTGACCTACTG |
| | FW: AACCATAGCCTTCACCTCCC |
| *Mlkl* | RV: GGATTGCCCTGAGTTGTTGC |
| | FW: AACCGCAGACAGTCTCTCCA |
| *18s* | FW: TCAAGAACGAAAGTCGGAGG |
| | RV: GGACATCTAAGGGCATCAC |
| **Human** | |
| *ACTA2* | FW: TTCAATGTCCCAGCCATGTA |
| | RV: GAAGGAATAGCCACGCTCAG |
| *COL1A1* | FW: CGGACGACCTGGTGAGAGA |
| | RV: CATTGTGTCCCCTAATGCCTT |
| *FN1* | FW: GTGTTGGGAATGGTCGTGGGGAATG |
| | RV: CCAATGCCACGGCCATAGCAGTAGC |
| *18S* | FW: AGCTATCAATCTGTCAATCCTGTC |
| | RV: GCTTAATTTGACTCAACACGGGA |

**Supplemental Table 3.** Primer sequence for qPCR analysis.

| Biological function | Abbreviation | Gene name |
|---|---|---|
| **Apoptosis** | ***Apaf1*** | Apoptotic Peptidase Activating Factor 1 |
| | ***Bax*** | Bcl-2-associated X protein |
| | ***Bcl2*** | B-cell lymphoma 2 |
| | ***Bcl2l1*** | (BCL2-like 1) |
| **Fatty acid oxidation** | ***Cpt1a*** | Carnitine palmitoyltransferase I |
| | ***Cpt2*** | Carnitine palmitoyltransferase II |
| | ***Ppara*** | Peroxisome proliferator-activated receptor alpha |
| | ***Ppargc1a*** | PPARG Coactivator 1 Alpha |
| | ***Acox1*** | Peroxisomal Acyl-CoA Oxidase 1 |
| | ***Acox2*** | Peroxisomal Acyl-CoA Oxidase 2 |
| **Fibrosis** | ***Acta2*** | alpha smooth muscle actin |
| | ***Cdh16*** | Cadherin 16 |
| | ***Col1a1*** | Collagen type I alpha 1 chain |
| | ***Col3a1*** | Collagen type III alpha 1 chain |

| | | |
|---|---|---|
| | ***Col4a1*** | Collagen type IV alpha 1 chain |
| | ***Fn1*** | Fibronectin |
| | ***Havrc1*** | Kidney Injury Molecule-1 |
| | ***Snai1*** | Snail Family Transcriptional Repressor 1 |
| | ***Tgfb1*** | Transforming Growth Factor Beta 1 |
| | ***Twist1*** | Twist Family BHLH Transcription Factor 1 |
| | ***Vim*** | Vimentin |
| **Glycolysis** | ***G6pc*** | Glucose-6-Phosphatase Catalytic Subunit |
| | ***Gapdh*** | Glyceraldehyde-3-Phosphate Dehydrogenase |
| | ***Hk1*** | Hexokinase 1 |
| | ***Ldh1*** | L-lactate dehydrogenase 1 |
| | ***Ldh2*** | L-lactate dehydrogenase 2 |
| | ***Pgk1*** | Phosphoglycerate Kinase 1 |
| | ***Pkfm*** | fosfofructoquinasa-1 |
| | ***Pkm*** | pyruvate kinase |
| | ***Slc2a1*** | Glucose transporter protein type 1 |
| **Inflammation** | ***Adgre1*** | F4/80 |
| | ***Agr1*** | Arginase-1 |
| | ***Cd86*** | CD86 Antigen |
| | ***IL1b*** | Interleukin 1 Beta |
| | ***IL6*** | Interleukin 6 |
| | ***Mrc1*** | Macrophage mannose receptor 1 (CD206) |
| | ***Nos2*** | Nitric Oxide Synthase 2 |
| | ***Tnf*** | Tumor Necrosis Factor |
| **Mitochondria** | ***Hspa9*** | Heat Shock Protein Family A (Hsp70) Member 9 |
| | ***Lrpprc*** | Leucine-rich PPR motif-containing protein |
| | ***Ndufv2*** | NADH:Ubiquinone Oxidoreductase Core Subunit V2 |
| | ***Sdha*** | Succinate Dehydrogenase Complex Flavoprotein Subunit A |
| | ***Tfam*** | Mitochondrial Transcription Factor 1 |

**Supplemental Table 4.** Genes analyzed with TaqMan probes.

| Antibody | Antibody Reference | Isotype control antibody | Isotype control antibody Reference |
|---|---|---|---|
| **APC anti-human CD45** | 368511 (Clone 2D1) | APC Rat IgG2b | 400611 (clone RTK4530) |
| **FITC anti-mouse F4/80** | 123107 (Clone BM8) | FITC Rat IgG2a | 400505 (Clone RTK2758) |
| **APC anti-mouse CD86** | 105113 (Clone PO3) | APC Rat IgG2b | 400611 (Clone RTK4530) |
| **PE anti-mouse CD206** | 141705 (Clone C068C2) | PE Rat IgG2a | 400507 (Clone RTK2758) |
| **APC/Cy7 anti-mouse CD326 (Ep-CAM)** | 118217 (Clone G8.8) | APC/Cy7 Rat IgG2a | 400523 (Clone RTK2758) |
| **PE anti-mouse CD24** | 101807 (Clone M1/69) | PE Rat IgG2b | 400607 (Clone RTK4530) |

**Supplemental Table 5.** Antibodies for flow cytometry analysis. All were from Biolegend (San Diego, CA).

| Antibody | Reference | Dilution |
|---|---|---|
| ***Primary antibodies*** | | |
| **anti-CPT1A** | Ab128568, Abcam | 1:1000 |
| **Biotinylated lotus tetragonolobus lectin (LTL)** | B-1325, Vector Laboratory | 1:1000 |
| **anti-β-F1-ATPase** | Clone 11/21-7 A8, 1: 1.000), kindly provided by Dr. Jose Manuel Cuezva (UAM, Madrid, Spain) (1) | 1:1000 |
| ***Fluorochrome-conjugated secondary antibodies*** | | |
| **Alexa Fluor 555 anti-Mouse** | A-21137, Thermo Scientific | 1:1000 |
| **Alexa Fluor 488 anti-Rabbit** | A-21206, Thermo Scientific | 1:1000 |
| **Streptavidin Alexa Fluor™ 555 Conjugate** | S32355, Thermo Scientific | 1:400 |

**Supplemental Table 6.** Antibodies for immunofluorescence.

| | **GFR<60<br>N=110** | **GFR≥60<br>N=576** | **P value** |
|---|---|---|---|
| **Age** | 71.2 (5.1) | 66.7 (6.0) | <0.001 |
| **Sex, women (%)** | 64 (58.2%) | 295 (51.2%) | 0.180 |
| **BMI, kg/$m^2$** | 29.9 (3.6) | 29.9 (3.5) | 0.942 |
| **Waist, cm** | 101.4 (11.1) | 100.8 (9.6) | 0.570 |
| **Smoking (%)** | | | |
| **Never** | 71 (64.6%) | 328 (56.9%) | 0.326 |
| **Current** | 14 (12.7%) | 94 (16.3%) | |
| **Former** | 25 (22.7) | 154 (26.7%) | |
| **Intervention group, (%)** | | | |
| **Control** | 35 (31.8%) | 200 (34.7%) | 0.067 |
| **MedDiet+EVOO** | 48 (43.6%) | 188 (32.6%) | |
| **MedDiet+nuts** | 27 (24.6%) | 188 (32.6%) | |
| **Diabetes, yes (%)** | 33 (30.0%) | 178 (30.9%) | 0.851 |
| **Hypertension, yes (%)** | 99 (90.0%) | 513 (89.1%) | 0.771 |
| **Dyslipidemia, yes (%)** | 75 (68.2%) | 433 (75.2%) | 0.125 |
| **Total cholesterol, mg/dl** | 216.1 (36.2) | 212.7 (36.6) | 0.369 |
| **Triglycerides, mg/dl** | 145.0 (66.7) | 133.4 (66.0) | 0.092 |
| **HDL, mg/dl** | 51.1 (11.3) | 51.4 (11.0) | 0.783 |
| **Systolic BP, mmHg** | 155.2 (19.7) | 149.9 (18.8) | 0.008 |
| **Diastolic BP, mmHg** | 83.6 (10.4) | 83.9 (9.7) | 0.707 |
| **Urinary creatinine,mg/dl** | 90.0 (52.5) | 95.0 (53.0) | 0.316 |
| **Urinary albumin, mg/L** | 24.4 (53.6) | 13.9 (34.3) | 0.008 |
| **eGFR (ml/min/1.73 $m^2$)** | 51.3 (7.5) | 80.9 (12.3) | <0.001 |

**Supplemental Table 7.** Characteristics of the participants by GFR levels in the PREDIMED study.

| | **GFR<60** | **GFR≥60** | **P value** |
|---|---|---|---|
| **Short-chained<br>(c2 to c7)** | 0.58 (0.40 to 0.77) | -0.11 (-0.19 to -0.03) | <0.001 |
| **Medium-chained<br>(c8 to c14)** | 0.25 (0.06 to 0.44) | -0.05 (-0.13 to 0.03 ) | 0.004 |
| **Long-chained<br>(c16 to c26)** | 0.03 (-0.16 to 0.21) | 0.00 (-0.08 to 0.07) | 0.782 |

**Supplemental Table 8.** Mean values (95% CI) of z-score standardized acyl-carnitine scores by GFR levels adjusted for age, sex, diabetes and albumin/creatinine ratio.

| | GFR<60 | GFR≥60 | P value |
|---|---|---|---|
| **Free carnitine** | 0.18 (-0.02 to 0.37) | -0.03 (-0.12 to 0.05) | 0.051 |
| **c2carnitine** | 0.28 (0.08 to 0.47) | -0.05 (-1.35 to 0.03) | 0.002 |
| **c3carnitine** | 0.09 (-1.01 to 0.29) | -0.02 (-0.10 to 0.07) | 0.323 |
| **c3dcch3carnitine** | 0.60 (0.41 to 0.79) | -0.11 (-0.19 to -0.04) | <0.001 |
| **c4carnitine** | 0.25 (0.06 to 0.44) | -0.05 (-0.13 to 0.03) | 0.006 |
| **c4ohcarnitine** | 0.33 (0.14 to 0.52) | -0.06 (-0.14 to 0.02) | <0.001 |
| **c5carnitine** | 0.52 (0.33 to 0.70) | -0.09 (-0.18 to -0.02) | <0.001 |
| **c51carnitine** | 0.41 (0.22 to 0.60) | -0.08 (-0.16 to 0.00) | <0.001 |
| **c5dccarnitine** | 0.70 (0.51 to 0.88) | -0.13 (-0.21 to -0.06) | <0.001 |
| **c6carnitine** | 0.13 (-0.6 to 0.32) | -0.02 (-0.11 to 0.06) | 0.145 |
| **c7carnitine** | 0.17 (-0.03 to 0.36) | -0.03 (-0.11 to 0.05) | 0.066 |
| **c8carnitine** | 0.20 (0.00 to 0.39) | -0.04 (-0.12 to -0.04) | 0.031 |
| **c9carnitine** | 0.10 (-0.09 to 0.29) | -0.02 (-0.10 to 0.06) | 0.285 |
| **c10carnitine** | 0.20 (0.01 to 0.39) | -0.04 (-0.12 to 0.04) | 0.025 |
| **c102carnitine** | 0.30 (0.11 to 0.49) | -0.06 (-0.14 to 0.02) | <0.001 |
| **c12carnitine** | 0.20 (0.01 to 0.39) | -0.04 (-0.12 to 0.04) | 0.024 |
| **c121carnitine** | 0.33 (0.14 to 0.52) | -0.06 (-0.14 to 0.02) | <0.001 |
| **c14carnitine** | 0.02 (-0.18 to 0.21) | -0.00 (-0.09 to 0.08) | 0.829 |
| **c141carnitine** | 0.28 (0.09 to 0.47) | -0.05 (-0.13 to 0.03) | 0.002 |
| **c142carnitine** | 0.13 (-0.06 to 0.33) | -0.03 (-0.11 to 0.06) | 0.139 |
| **c16carnitine** | -0.07 (-0.27 to 0.12) | 0.01 (-0.07 to 0.10) | 0.418 |
| **c18carnitine** | -0.05 (-0.24 to 0.14) | 0.01 (-0.07 to 0.09) | 0.570 |
| **c181carnitine** | 0.19 (0.01 to 0.38) | -0.04 (-0.12 to 0.41) | 0.025 |
| **c181ohcarnitine** | 0.21 (0.02 to 0.40) | -0.04 (-0.12 to 0.04) | 0.021 |
| **c182carnitine** | -0.06 (-0.25 to 0.13) | 0.01 (-0.07 to 0.09) | 0.508 |
| **c20carnitine** | -0.11 (-0.30 to 0.09) | 0.02 (-0.06 to 0.10) | 0.232 |
| **c204carnitine** | -0.03 (-0.21 to 0.16) | 0.01 (-0.07 to 0.08) | 0.748 |
| **c26carnitine** | -0.08 (-0.27 to 0.11) | 0.02 (-0.07 to 0.10) | 0.362 |

**Supplemental Table 9.** Adjusted (for age, sex, diabetes and albumin/creatinine ratio) mean values (95% CI) of z-score individual standardized acyl-carnitines (named according the HMDB) by GFR levels. HMDB: Human Metabolome Database

| | Primary Cohort |
|---|---|
| **Subjects (n)** | N=433 |
| **GFR (ml/min/1.73 m2 by CKD-EPI)** | 67.3 (26.71) |
| **Gender (%Female)** | 160 (36.95%) |
| **Age** | 61.39 (12.49%) |
| **Race** | |
| **Asian (n)** | 7 (1.62%) |
| **Caucasian (n)** | 290 (66.97%) |
| **African American (n)** | 1 (0.23%) |
| **Hispanic (n)** | 77 (17.78%) |
| **Multi-racial (n)** | 10 (2.31%) |
| **Diabetes** | 164 (37.88%) |
| **Hypertension** | 316 (72.98%) |

| | |
|---|---|
| **Dipstick Protein (0=neg, 1=trace, 2=30, 3=100, 4=300, 5>300)** | 1.06 (1.56) |
| **BMI (kg/m2)** | 30.79 (7.88) |
| **HgbA1c** | 6.53 (1.25) |
| **Serum glucose (mg/dl)** | 133.08 (64.48) |
| **BP systolic (mmHg)** | 136.88 (21.32) |
| **Serum-alb (gldl)** | 4.19 (2.87) |
| **Glomeruli: Hypoperfused: 0-3** | 0.86 (0.71) |
| **Glomeruli: Wall Thickening: 0-3** | 0.24 (0.59) |
| **Glomeruli: Mesangial Matrix: 0-3** | 0.47 (0.82) |
| **Glomeruli: Mesangial Cellularity: 0-3** | 0.38 (0.75) |
| **Glomeruli: KW Nodules: 0-1** | 0.04 (0.2) |
| **Glomeruli: Pericapsular Fibrosis: 0-2** | 0.78 (0.7) |
| **Glomeruli: Globally Schlerotic %** | 14.13 (19.61) |
| **Tubules: % Atrophy** | 14.37 (23.01) |
| **Tubules: % Acute Tubular Injury** | 3.26 (7.94) |
| **Tubules: Reabsorption: 0-3** | 0.41 (0.62) |
| **Interstitium: % Fibrosis** | 14.34 (21.68) |
| **Interstitium: Lymphocytic Infiltrate: 0-3** | 1.09 (0.87) |
| **Interstitium: Plasmacytic Infiltrate: 0-3** | 0.43 (0.65) |
| **Interstitium: Eosinophils: 0-3** | 0.27 (0.51) |
| **Vessels: Medial Thickening: 0-3** | 0.15 (0.44) |
| **Vessels: Intimal Fibrosis: 0-3** | 1.52 (0.85) |
| **Vessels: Arteriolar Hyalinosis: 0-3** | 0.56 (0.78) |

**Supplemental Table 10.** Demographic and clinical information of the CKD cohort. Data are mean (SD) or n (%), which are reported as counts for discrete variables and reported as mean and SD for continuous variables.

**Supplemental Methods**

*Unilateral ureteral obstruction (UUO).* UUO surgery procedure was performed as previously described (2). Briefly, mice were anesthetized with isoflurane (3-5% for induction and 1-3% for maintenance) and divided into two experimental groups: the UUO group and the sham operation group. In the UUO group, mice were shaved on the left side of the abdomen, a vertical incision was made through the skin with a scalpel and the skin was retracted. A second incision was made through the peritoneum to expose the kidney. The left ureter was ligated twice 15 mm below the renal pelvis with surgical silk and the ureter was then severed between the two ligatures. Then, the ligated kidney was placed gently back into its correct anatomical position and sterile saline was added to replenish loss of fluid. The incisions were sutured and mice were individually caged. The sham operation was performed in a similar manner, but without ureteral ligation. Buprenorphine was used as an analgesic. A first dose was administered 30 minutes before surgery and then every 12 hours for 72 hours, at a dose of 0.05 mg / kg subcutaneously. In this model, renal blood flow and glomerular filtration rate become significantly reduced within 24 h and interstitial inflammation (peak at 2–3 days), tubular dilation, tubular atrophy and fibrosis are evident after 7 days. The obstructed kidney reaches maximal dysfunction around 2 weeks after the procedure. Mice were sacrificed by $CO_2$ overdose and control and obstructed kidney and blood samples were harvested after perfusion with PBS at 3 and 7 days after UUO.

*Folic acid-induced nephropathy (FAN).* In this model, kidney fibrosis was induced by intraperitoneal (i.p) injection with 250 mg folic acid (Sigma-Aldrich, St. Louis, MO) per kg body weight dissolved in 0.3 M sodium bicarbonate (vehicle) as previously described (3). Control animals received 0.3 ml of vehicle (i.p). Mice were sacrificed by $CO_2$ overdose and kidneys and blood samples were harvested after perfusion with PBS after 15 days of FA administration.

*Adenine-induced renal failure (ADN).* Adenine (Sigma-Aldrich, St. Louis, MO) was administered daily to mice at 50 mg/kg body weight in 0.5% carboxymethyl cellulose (CMC) (Wako Pure Chemical Industries Ltd., Osaka, Japan) by oral gavage for 28 days as previously described (4). Control animals received daily 0.5% CMC administered by oral gavage for the same period of time. Mice were sacrificed by $CO_2$ overdose and kidneys and blood samples were harvested after perfusion with PBS after 28 days of the first adenine dose administration.

*Measurements of oxygen consumption rate.* HKC-8 cells were seeded in a p60 plate and microRNA transfection or adenovirus-mediated CPT1A overexpression was performed (as shown in the transfection procedure and adenovirus-mediated overexpression sections, respectively) when they reached a confluence of 70% and 48 hours later, HKC-8 cells were treated with 10 ng/ml TGF-β1 for 48 h. In the case of primary kidney epithelial cells, they were seeded in a p60 plate and when they reached a confluence of 70%, they were treated with 10 ng/ml TGF-β1 for 48 h. In all cases, cells were then seeded at $2 \times 10^4$ cells per well in a Seahorse Bioscience XFe24 cell culture microplate (Seahorse Bioscience, North Billerica, MA). After cell adherence, growth medium was replaced with substrate-limited medium, Dulbecco's Modified Eagle Medium (DMEM) supplemented with 0.5 mM Glucose and 1 mM Glutamate. One hour before the assay measurement, cells were incubated with Krebs-Henseleit Buffer (KHB) assay medium supplemented with 0.2% carnitine at 37ºC without $CO_2$. Fifteen minutes before the assay, the CPT1 inhibitor Etomoxir (Eto) (Sigma-Aldrich, St. Louis, MO) 400 µM was added to the corresponding wells and cells were incubated at 37ºC without CO2. Finally, just before starting the assay, BSA or 200 µM Palmitate-BSA FAO Substrate (Agilent Technology, Santa Clara, CA, USA) was added.

Immediately, XF Cell Mito Stress Test was performed in a Seahorse XFe24 energy analyzer by adding sequentially during the assay several modulators of mitochondrial function: 1 μM oligomycin (Sigma-Aldrich, St. Louis, MO), 3 μM cyanide 4-(trifluoromethoxy) phenylhydrazone (FCCP) (Sigma-Aldrich, St. Louis, MO) and 1 μM antimycin/rotenone (Sigma-Aldrich, St. Louis, MO), Values were normalized for total protein content. Basal mitochondrial respiration, ATP-linked respiration, proton leak (non-ATP–linked oxygen consumption), maximal respiration, non-mitochondrial respiration and reserve respiratory capacity were determined as described (5). In the case of the reversion experiments (Fig.7) OCR and ATP production rate were determined in a Seahorse XFe96 analyzer by using the Agilent Seahorse XF Real-Time ATP Rate Assay in the latter case. Cells were seeded at $10^4$ cells per well. The injection of 1 μM oligomycin and 1 μM antimycin/rotenone enabled the calculation of mitochondrial and glycolytic ATP production by using Agilent ATP Assay Report Generator. Four wells were used for each experimental group.

*Histological and immunohistochemical analysis.* Fibrosis was quantified in Sirius red-stained sections in order to detect collagen fibers. The area of interstitial fibrosis was identified, after excluding the vessel area from the region of interest, as the ratio of interstitial fibrosis or collagen deposition to total tissue area and expressed as %FA (fibrotic area). Tubular atrophy and tubular dilation were evaluated as previously described in PAS-stained sections (6). Renal T lymphocyte and macrophage infiltration was quantified by CD3 and F4/80 stained area versus total analyzed area, respectively. For each kidney, 10–15 fields were analyzed with a 40X objective lens under transmitted light microscopy by using a digital camera (Nikon D3) connected to a Nikon's Eclipse TE2000-U light microscope (Nikon Instruments Europe B.V., Badhoevedorp, The Netherlands). Intensity measurements of Sirius red, CD3 and F4/80 staining area were performed blindly in an automated mode using the ImageJ 1.48 software (http://rsb.info.nih.gov/ij). Furthermore, qualitative evaluation of the tissue was performed by a renal pathologist in a blinded fashion.

*Production of adenovirus-mediated CPT1A overexpression*. In consisted of two rounds of amplification. In the first amplification round, HEK293A cells in a 100 $cm^2$ dish at 70% confluence growing in complete medium supplemented with 5% of FBS (infection media) were infected with 50 µl of the adenovirus stock, while in the second amplification round, 10 X 100 $cm^2$ dishes of 70% confluent HEK293A cells /virus type were infected with 400 µl of the first amplification of the adenovirus. After 20 h, cells from all dishes were harvested together by scraping and centrifuged (1,000 rpm for 5 min at 4ºC). The cell pellet was disrupted by 3 cycles of freeze and thaw using liquid $N_2$ and resuspended in 5 mL of infection media. Cell debris was discarded by centrifugation at 3,000 rpm for 10 min at 4ºC and supernatant containing adenoviruses was stored in 500 µl aliquots to be used throughout the experiments. Titration was performed by using the Adeno-X™ Rapid Titer Kit (Clontech Laboratories, Palo Alto, CA) according to the manufacturer's instructions, which is based on the antibody specific detection of the adenoviral hexon protein by peroxidase (HRP).

*Data in clinical studies for acyl-carnitines*: Means and standard deviations (SD) were used to describe continuous variables and percentage to describe categorical values. Metabolite measures were normalized using a logarithmic transformation and scaled to multiples of SD using z-score standardization. Three scores were created according to the number of carbons: short (2-7), medium (8-14) and long (16-26) acyl-carnitine scores. The sum of the log-transformed values of each carnitine was computed to calculate these scores and a z-score standardization was also applied. Chi-squared and Student´s t-test were used to compare categorical and quantitative variables, respectively. Univariate and multivariable linear regression models were conducted to assess the association between the glomerular filtration rate (GFR) and both the acyl-carnitine scores and individual metabolites. The covariates used in the multivariable models were age, sex, type-2 diabetes and albumin/creatinine ratio.

*Data in clinical studies for CPT1A expression:* ANOVA test was performed to assess the significance across different disease groups and cor.test() function in R was used to get the Pearson's correlation and the corresponding P-values.
